# Development of a microRNA clock for gestational age in the general population

**DOI:** 10.64898/2026.05.22.727152

**Authors:** Tim Finke, Midas M. Kuilman, Frédérique White, Janine F. Felix, Vid Prijatelj, Isabel K. Schuurmans, Henning Tiemeier, Neeltje van Haren, Luigi Bouchard, Pierre-Étienne Jacques, Marie-France Hivert, Mohsen Ghanbari, Alexander Neumann, Charlotte A. M. Cecil

**Author notes:** Shared first author (Tim Finke, Midas M. Kuilman). Shared senior author (Alexander Neumann, Charlotte A. M. Cecil).

## Abstract

**Introduction:** DNA methylation (DNAm) has shown promise as a biological marker of gestational age (GA). Here, we aimed to characterize how plasma circulating miRNAs – another epigenetic mechanism regulating gene expression – associate with GA at birth, and to construct a miRNA-based GA clock (miRClock-GA).

**Methods:** We leveraged 2083 umbilical cord plasma-derived circulating miRNAs from Generation R (N=1695). First, we performed linear regressions to identify miRNome-wide significant miRNAs associated with GA. Second, we applied elastic net regression to construct miRClock-GA. These steps were validated in Gen3G (N=213). Finally, we computed age acceleration (miRClock-AA) and evaluated association of miRClock-GA and miRClock-AA with child developmental outcomes up to 17 years of age, including comparisons with DNAmClocks.

**Findings:** We identified 123 miRNAs associated with GA, with miR-150-5p showing the strongest positive association ( *B*=0.244, SE=0.036, P=2.3e-11) and miR-373-3p the strongest negative association ( *B*=-0.255, SE=0.065, P=8.6e-5). MiRClock-GA correlated consistently with GA in Generation R train (r_*range*_=0.62-0.72) and test sets (r_*range*_=0.45-0.52) and in the independent validation cohort (r_*Gen*3*G*_ =0.33). Correlations between miRClock-GA and DNAmClocks were weak to moderate (r_*range*_=0.28-0.42). MiRClock-AA explained significant variance in birthweight and childhood BMI beyond clinical GA.

**Conclusions:** This study reveals widespread associations between circulating miRNAs and GA, supports miRClock-GA as a consistent, well-performing biological marker of GA, with miRClock-AA predicting birthweight and childhood BMI beyond GA itself. Our findings provide a broader perspective on the potential utility of miRNAs as early markers of development.

**Funding:** E.U. Horizon Europe Research and Innovation Programme (FAMILY,No.101057529); European Research Council (TEMPO,No.101039672). Full funding in Acknowledgements.

**Research in context:** *Evidence before this study:* Epigenetic clocks have emerged as powerful tools for assessing individual differences in biological ageing. Existing epigenetic clocks are typically constructed using DNA methylation (DNAm) data and are developed to estimate whether an individual’s epigenetic age deviates relative to their chronological age. In adults, higher predicted biological epigenetic age than chronological age (accelerated epigenetic ageing) is an established indicator of mortality risk and other age-related morbidities. MicroRNAs are another epigenetic mechanism associated with chronological age and age-related phenotypes across species such as *C. elegans*, mice, primates and humans. Recently, a plasma-derived, cell-free circulating miRNA clock was developed in a sample of older European adults, with accelerated miRNA age predicting biological age-related conditions such as frailty and multi-system blood biomarkers. However, miRNA-based estimations of biological gestational age at birth have not been previously done and their relationship to child health outcomes are unknown.

*Added value of this study:* To gain a broader perspective on the role of epigenetic markers in biological gestational age at birth and the potential utility of miRNAs as early markers of (altered) development, we leveraged cord blood plasma circulating miRNA data collected at birth in a large population-based cohort. First, we identified 123 miRNAs associated with gestational age. Second, we used this information to construct a miRNA-based epigenetic clock for gestational age (miRClock-GA) and validated this clock in an independent cohort. MiRClock-GA showed moderate correlations with both gestational age and existing DNAm-based gestational clocks. Third, we found that miRClock-GA correlated cross-sectionally with birthweight, as well as prospectively with several adaptive, behavioural, cognitive, and growth outcomes measured up to age 17y, after taking into account maternal influences such as maternal education level, smoking, pre-pregnancy BMI, and age. Accelerated ageing of miRClock-GA (miRClock-AA) explained variance in birthweight and childhood BMI beyond clinical GA, after accounting for maternal influences.

*Implications of all available evidence:* Our study provides the first large-scale evidence that circulating miRNAs in cord blood plasma show widespread associations with gestational age at birth and can be used to derive a robust biological marker of GA (MiRClock-GA), which explains a substantial proportion of the variation in clinical GA. MiRClock-AA provides information beyond GA in some child outcomes, such as BMI. These findings extend adult research, supporting the potential of miRNAs as promising markers for early risk assessment and health monitoring in a developmental context.

## Introduction

Epigenetic clocks have emerged as powerful tools for assessing individual differences in biological ageing.^1,2^ Typically constructed using DNA methylation (DNAm) data, these clocks estimate whether an individual is ageing relatively fast (acceleration) or slow (deceleration) compared to peers of the same chronological age. While DNAm-based accelerated epigenetic age has been consistently linked to increased mortality risk and age-related morbidities in adulthood,^3–6^ its applicability as a risk predictor in early life is less established.

A handful of DNAm-based clocks to date have been specifically trained to estimate gestational age (GA) at birth, with implications for developmental and health outcomes later in life.^7–10^ However, these gestational clocks show inconsistencies in the direction and strength of associations with prenatal risk factors (e.g. maternal smoking, pre-eclampsia, gestational diabetes)^11,12^ and childhood outcomes (e.g. blood pressure, carotid intima-media thickness, ADHD symptoms).^13,14^

While DNAm has been the predominant focus of epigenetic clock research, circulating microRNAs (miRNAs) have recently emerged as promising markers of age-related processes and may thus complement existing clocks based on DNAm data.^15^ MiRNAs are small, non-coding RNAs of 20 - 25 nucleotides, which are involved in post-transcription gene regulation in response to both internal (e.g. genetic) and external (e.g. environmental) inputs.^16^ When released from cells into extracellular environments (e.g. plasma, serum), circulating miRNAs are relatively stable,^17^ enabling inferences beyond acute states. Their associations with chronological age and relevance to age-related phenotypes has been increasingly demonstrated across species such as *C. elegans*, mice, primates and humans.^18^ For example, a plasma-derived, cell-free circulating miRNA clock was recently developed in a sample of older European adults, with accelerated miRNA age predicting biological age-related conditions such as frailty and blood biomarkers.^19^

Within a developmental context, differential miRNA expression in cord blood has been associated with prenatal exposures and gestational factors linked to offspring development and health, including maternal age, tobacco smoking, BMI, gestational hypertension, preeclampsia, and fetal growth restriction.^20–23^ However, no studies have yet investigated the extent to which cord blood plasma circulating miRNA profiles associate with GA itself, whether it is possible to use this information to construct a miRNA-based GA clock (miRClock-GA), and how predictive such a clock is of later developmental outcomes. Addressing these gaps is critical to gain a broader perspective on the role of different epigenetic markers in biological age(ing) and the potential utility of miRNAs as early markers of (altered) development.

To address these gaps, this preregistered study (https://osf.io/7yb4m/) leveraged data of 2083 miRNAs from 1695 children from the general Dutch population, with three key aims: (1) to conduct a miRNome-wide association analysis in order to identify plasma circulating miRNAs associated with GA at birth; (2) to construct and validate a plasma circulating miRNA-based GA clock (miRClock-GA); and (3) to map prospective associations between miRClock-GA and developmental outcomes at later ages, including comparisons with existing DNAm-based GA clocks. Across the first two steps, we evaluated the robustness and generalizability of our findings in an independent cohort of 213 Canadian children. We additionally conducted downstream *in silico* analyses to explore the functional significance of the top miRNAs associated with GA and tested overlap with previously-identified age-associated miRNAs in older life.

## Method

### Study Settings

This study included participants from the population-based prospective Generation R Study (GenR).^24^ GenR investigates early environmental and genetic factors leading to normal and abnormal growth, development and health from fetal life until adulthood. A total of 9778 pregnant women residing in Rotterdam, with delivery dates between April 2002 and January 2006 were enrolled in the study. Out of this cohort, a subset of 1710 participants was selected for miRNA sequencing of umbilical cord blood-derived plasma, based on availability of (i) DNAm profiles already estimated at birth in GenR, (ii) follow-up data in later waves of data collection, and (iii) not having a sibling that was already in the subset. After data normalisation and cleaning, the analyses include a subset of 1695 participants with miRNA data (**Figure 1**).

**Figure 1.**
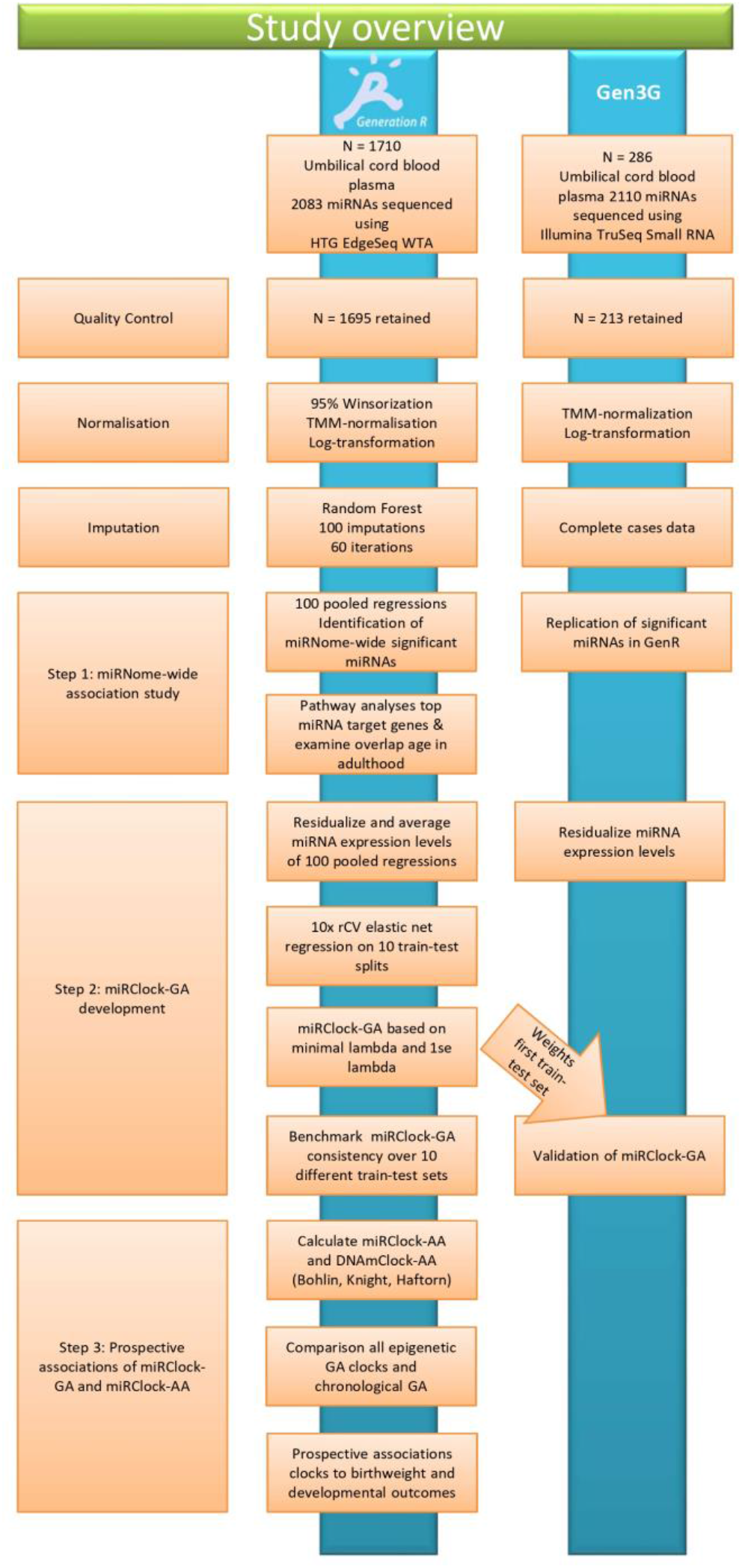
Study overview.

Additionally, we included 213 participants from the prospective prebirth cohort Genetics of Glucose regulation in Gestation and Growth (Gen3G) in Canada.^25,26^ Pregnant women were recruited between 2010 and 2013 during their first trimester of pregnancy. Biological samples, including umbilical cord blood at delivery, were obtained for DNAm and miRNA profiling. Unrelated participants with both DNAm and miRNA sequencing data available from cord blood were included for replication analyses of the GenR findings.

### Measures

A full description of miRNA expression profiling and normalization, DNA methylation-derived data, and other variables used is present in the supplementary materials.

#### MiRNAs expression profiling and normalization in Generation R

Plasma circulating miRNAs from umbilical cord blood were profiled in 1710 newborns using the HTG EdgeSeq miRNA Whole Transcriptome Assay^27^ and sequenced on an Illumina NextSeq 2000.^28^ Reads were quality-checked and conservatively aligned using Bowtie2^29^ to a miRBase v20–based reference.^30^ After quality control, 1695 samples with expression data for 2083 miRNAs were retained. Data were first adjusted for total reads within each sample and subsequently TMM-normalized,^31^ log-counts-per-million (log-CPM) transformed, and 95% winsorized to limit false positive inflation, e.g. by limiting the influence of outliers due to low expression.^32,33^

#### Gestational age estimation in Generation R

Pregnant mothers received an ultrasound in the first trimester of pregnancy at our research center^34^. If mothers could reliably recall the first day of their last menstrual period (LMP), and had a regular menstrual cycle of 28 (+/-4) days, a second clinical estimate of GA was calculated from LMP and used in sensitivity analyses.

#### Developmental outcomes from birth to adolescence

To assess the association of miRClock-GA with later child development varying from 3 months to 17 years of age, we focused on 51 variables categorized along five developmental domains, encompassing (i) adaptive, (ii) cognitive, (iii) physical, (iv) social-emotional, and (v) communicative development.

#### Covariates

Covariates known to associate with health and developmental outcomes, as well as with GA, were included, namely self-reported maternal education (low/primary education, middle/secondary education, higher education), prenatal smoking (none; yes, until pregnancy was known; yes, continued during pregnancy), self-reported maternal pre-pregnancy BMI (kg/m^2^), and maternal age at intake. Technical covariates included the initial total sample miRNA concentrations prior to dilution in the lab, miRNA-sequencing plate, haemolysis levels, and blood cell counts. Haemolysis was assessed *ad oculos* in the lab using a four-level scale. For the analyses featuring DNAm-based clocks, we also included methylation array plate as a measure of batch effects in the DNAm data.

#### Validation cohort

For the Gen3G cohort, small RNA libraries were prepared from umbilical cord blood plasma using the TruSeq Small RNA Sample Prep Kit (Illumina, BC, Canada, catalog #RS-200-0012) following a low-input protocol and sequenced on an Illumina NovaSeq 6000 platform (100 cycles, single-end). Reads were processed using the exceRpt pipeline,^35^ with adapter trimming, contaminant removal, alignment to the human genome (GRCh38), and the transcriptome (miRBase v21^30^) using STAR, and miRNA quantification. After quality control, 213 samples with expression data for 2125 miRNAs were retained. Due to the number of miRNAs with low expression values, miRNAs were filtered for at least 5 CPM in at least 50% of samples. The remaining 680 miRNAs were normalized following the same procedure as GenR (total reads adjustment, TMM normalization and log-CPM transformation) and no winsorization was applied. Of these 680 miRNAs, 560 were present in GenR data as well.

In Gen3G, GA at birth was calculated as the number of weeks and days between LMP and delivery. LMP was initially determined from maternal report at the first trimester visit and subsequently confirmed by first trimester ultrasound. When discrepancies exceeded 5 days between reported- and ultrasound-derived dates, the ultrasound-based LMP was used for GA calculation. Covariates were comparable to those in GenR. Blood cell type proportions were estimated using the same method as in GenR.^36^ Haemolysis level was assessed using a visual analogue scale.^37^

### Statistical Analyses

#### Missing values

In GenR, missing values for DNAmClock-GA, GA, developmental outcomes, technical variables and other covariates were imputed using the R-package *mice*^38^, applying a random forest imputation with 100 imputations and 60 iterations. All outcome and covariate variables except for miRNA expression values were used as predictors in the imputation model. Additionally, we included country of origin and GA ultrasound measurement method in the imputation.

#### Step 1. MiRNome-wide association study for gestational age

To identify miRNAs associated with GA, we first performed linear regression analyses in the full dataset using GA as the outcome and expression levels of individual miRNAs as the predictor (each miRNA modelled individually), adjusting for sex, haemolysis, miRNA-sequencing plate, initial total sample miRNA concentrations prior to dilution, and blood cell counts as estimated from DNAm data. We applied Benjamini-Hochberg false discovery rate (FDR)-correction to control for multiple testing.^39^ As secondary analyses, linear regression analyses were re-run including an interaction term between miRNA expression level and sex. GA-associated miRNAs were then further characterized to (i) test their overlap with previously identified miRNAs associated with chronological age in adulthood,^19^ (ii) test whether they also associate with clinical GA at birth in the independent Gen3G cohort, and (iii) explore associated miRNAs’ experimentally validated target genes derived from miRDB^40^ and miRTarBase^41^ through assessing their genetic associations, and investigating their functional relevance using pathway enrichment analysis.

#### Step 2. Development of a gestational age miRClock (miRClock-GA)

Our approach followed the methodology previously applied to derive a miRClock from chronological age in adult samples.^15,19^ Elastic net models were run on the residuals of 2083 miRNA expression levels after adjustment for the same covariates used in the miRNome-wide association analysis (Step 1). These residuals were averaged across the 100 imputed datasets. We split our sample 60:40 randomly in a training set of 1017 and test set of 678 participants to evaluate the performance of the elastic net model. We repeated the train/test split ten times to assess consistency of the model across different train/test allociations. Subsequently, we conducted the elastic net regression model on the training set with clinical GA as the outcome, using the R-package *glmnet*.^42,43^ Elastic net models combine Lasso and Ridge regression, selecting the most informative miRNAs for building miRClock-GA. We optimized the regularization parameters for the elastic net model using 10-fold cross-validation to find the optimal combination of lambda and alpha to minimize the mean squared error using the cva.glmnet function from *glmnetUtils*.^44^ We also present a more parsimonious model in the supplementary materials with the highest shrinkage, which still falls within 1 standard error of the lowest mean squared error (1SE rule^45^). The coefficients from the selection were used to calculate miRClock-GA in the hold-out test set by summing the coefficients weighted by their effect size. This resulted in a measure of epigenetically predicted GA (i.e. miRClock-GA). From this, we derived a measure of age acceleration (miRClock-AA) by regressing miRClock-GA onto clinical GA, where positive values indicate being epigenetically “older” than one’s chronological age. Unless explicitly mentioned otherwise, we reported on the model applying minimal lambda for train-test split A (**Figure 1**), corresponding to the first train-test split of the 10x repeated CV models. The weights assigned to this train-test split were shared with Gen3G in order to apply them on the available miRNAs in Gen3G in the computation of miRClock-GA in the validation set.

Pearson correlations were applied to assess correlations between clinical GA (from clinical assessment) versus miRClock-GA within both the training and test sets, as well as the validation set in Gen3G, in order to assess the performance of miRClock-GA in predicting clinical GA. To account for method-specific variations in GA estimation,^13^ we performed a sensitivity analysis in a high-confidence GenR subsample to assess the correlation between miRClock-GA and clinical GA as measured through LMP as well as through ultrasound.

#### Step 3. Associations of miRClock-GA and miRClock-AA with developmental outcomes and comparison with DNAmClocks

Within the GenR test set, we examined to what extent miRClock-GA correlates with existing DNAmClock-GA (Bohlin, Knight, Haftorn) through Pearson correlations, including both measures of estimated GA and age acceleration per clock.

Next in GenR, we tested whether miRClock at birth was associated with a range of health and developmental outcomes from birth to adolescence, following a strategy previously applied to adult data.^46^ Specifically, we ran step-wise linear regression models (one per outcome) to calculate the additional explained variance (*R*^2^) between the following models: *Model 1* containing only covariates as predictors (same as those used in Step 1 and 2); *Model 2* containing covariates + clinical GA; and *Model 3* containing covariates + clinical GA + miRClock-AA. As an additional model (*Model 4*), we repeated these analyses with incorporation of an extended set of covariates, including maternal age, education level, smoking, and BMI in order to determine to what degree observed associations between miRClock-AA and outcomes are independent from these factors. To adjust for multiple testing of the 51 outcomes investigated, we applied Galwey correction for the number of effective tests (P<0.05 / 38).^47^ We first focused on associations with birthweight as a positive control, given that (1) it is assessed at the same time point as GA and the cord blood sampling; and (2) it is known to strongly associate with both clinical GA and epigenetically-estimated GA based on existing DNAmClocks. Next, we extended analyses to the other, prospective, health and developmental outcomes. We repeated these analyses using DNAmClock-AA (Bohlin, Knight, Haftorn) as a predictor to compare their pattern of associations with those observed using miRClock-AA.

## Results

### Characteristics of study participants

**Table 1.** Descriptive characteristics of Generation R and Gen3G.

| Variable | Generation R |  |  | Gen3G |  |  |
| --- | --- | --- | --- | --- | --- | --- |
|  | N | Mean | SD | N | Mean | SD |
| N females | 872 (51.4%) |  |  | 100 (46.9%) |  |  |
| Gestational age (weeks) | 1695 | 40.1 | 1.5 | 213 | 39.6 | 1.0 |
| Birth weight (g) | 1695 | 3545 | 493 | 213 | 3431 | 442 |
| Maternal age | 1695 | 31.8 | 4.1 |  |  |  |
| Maternal BMI | 1692 | 24.0 | 3.9 |  |  |  |
| Maternal education level |  |  |  |  |  |  |
| High | 1103 (65.1%) |  |  |  |  |  |
| Middle | 522 (30.8%) |  |  |  |  |  |
| Low | 43 (2.5%) |  |  |  |  |  |
| Maternal smoking |  |  |  |  |  |  |
| No | 1231 (72.6%) |  |  |  |  |  |
| Yes, in the past |  |  |  |  |  |  |
| Yes, until pregnancy was known | 143 (8.4%) |  |  |  |  |  |
| Yes, continued during pregnancy | 193 (11.4%) |  |  |  |  |  |
| Haemolysis |  |  |  |  |  |  |
| None | 1546 (91.2%) |  |  | 124 (58.2%) |  |  |
| A little | 111 (6.5%) |  |  | 59 (27.7%) |  |  |
| Some | 34 (2%) |  |  | 19 (15.4%) |  |  |
| Most | 4 (0.2%) |  |  | 8 (6.5%) |  |  |
| Cell type proportions |  |  |  |  |  |  |
| Bcell | 1622 | 0.057 | 0.023 | 213 | 0.053 | 0.028 |
| CD4T | 1622 | 0.124 | 0.059 | 213 | 0.169 | 0.053 |
| CD8T | 1622 | 0.043 | 0.032 | 213 | 0.031 | 0.022 |
| Granulocytes | 1622 | 0.514 | 0.099 | 213 | 0.548 | 0.105 |
| Monocytes | 1622 | 0.089 | 0.028 | 213 | 0.088 | 0.031 |
| NK | 1622 | 0.068 | 0.032 | 213 | 0.052 | 0.031 |
| nRBC | 1622 | 0.124 | 0.080 | 213 | 0.085 | 0.078 |

Descriptive characteristics of Generation R (N=1695) and Gen3G (N=213), including variables applied in Step 1, Step 2, and the maternal covariates applied in Step 3. A full overview of descriptive characteristics, including the health- and developmental outcomes at later ages applied in Step 3, is present in the supplementary materials. These descriptive characteristics are prior to imputation in the Generation R-sample. In Gen3G, missing smoking data was imputed for two participants.

#### MiRNome-wide association study of gestational age

Our analyses identified 123 circulating miRNAs exhibiting differential expression levels in relation to GA after FDR correction for multiple testing (**Figure 2A**). Among these, 106 (86%) miRNAs were upregulated (top 10 shown in **Figure 2B**), and 17 (14%) were downregulated (top 10 shown in **Figure 2C**). The top positively associated miRNA was miR-150-5p ( *B*=0.244, SE=0.036, P=2.3e-11, **Figure 3A & B**) and the top negatively associated miRNA was miR-373-3p ( *B*=-0.255, SE=0.065, P=8.7e-5, **Figure 3C & D**). Analyses including an interaction of each miRNA with sex did not yield significant interaction effects, suggesting that sex does not significantly moderate associations between miRNAs and gestational age. We found no overlap between our findings and miRNAs associated with chronological age in adults from the Rotterdam Study.^19^

**Figure 2.**
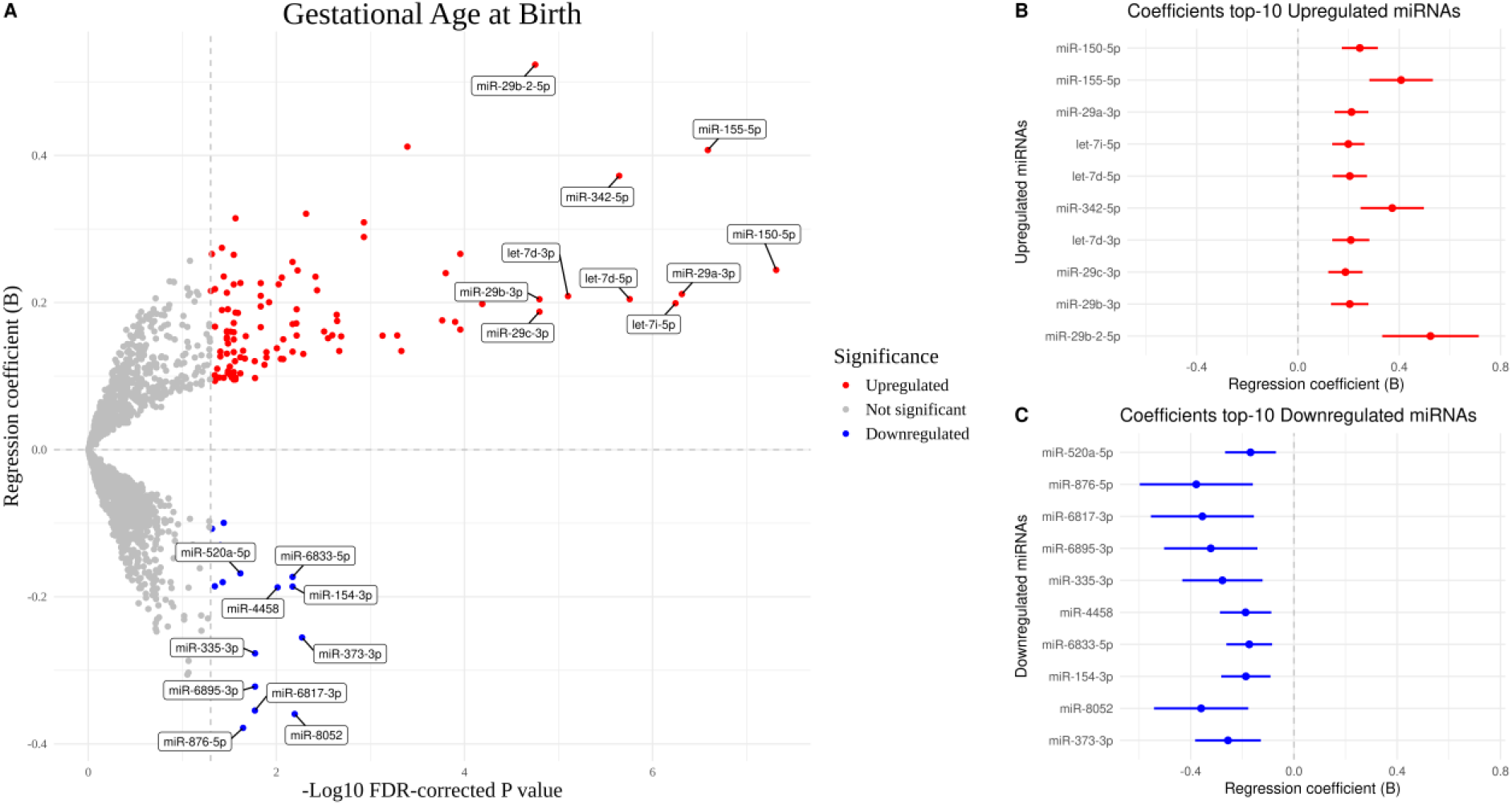
MiRNome-wide association study of gestational age. (A) Volcano plot of miRNAs associated with GA, with colour denoting associations that survive multiple testing correction; (B) the top 10 upregulated and (C) top 10 downregulated miRNAs with GA, including coefficients and their 95% confidence interval. Coefficients correspond to the amount of GA weeks changed for every 1 SD of change in miRNA expression levels.

**Figure 3.**
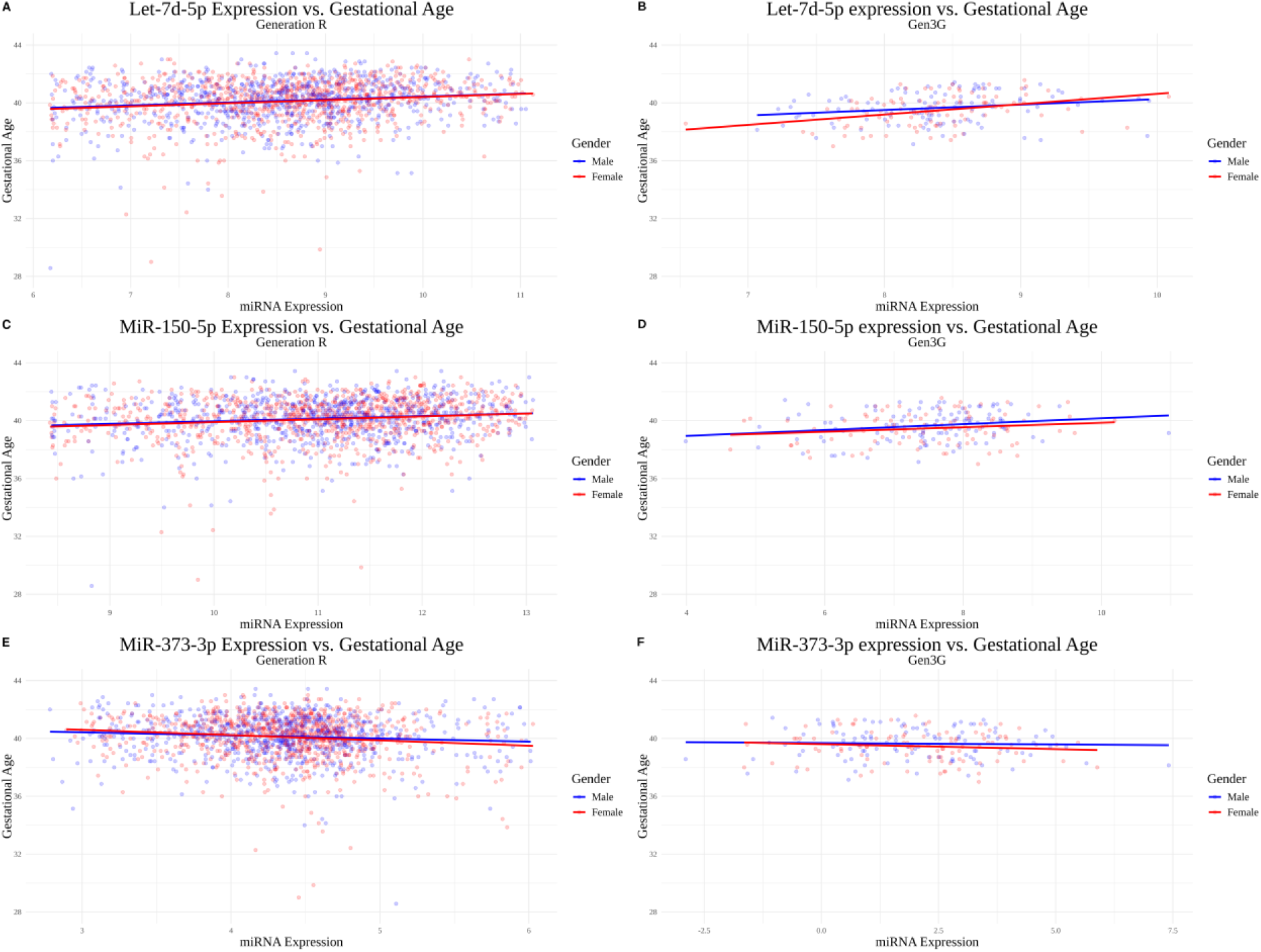
Top associated miRNAs miR-150-5p, miR-373-3p, and let-7d-5p for both Generation R (A, C, E) and Gen3G (B, D, F), coloured by sex. The lines portray the relation between miRNA expression and gestational age prior to correction for covariates for males and females separately.

We conducted downstream analyses to explore the functional significance of the top ten miRNAs (based on smallest P-values; **Figure 4A**). Notably, these miRNAs are all correlated positively with GA. The number of experimentally validated target genes for each miRNA ranged from 0 to 30 (**Figure 4A**). To further assess the biological significance of the top 10 miRNAs, genetic variants annotated to their experimentally validated target genes were extracted and linked to GA using a previous GWAS.^48^ After adjusting for multiple testing based on the number of unique target genes, several genes showed significant associated variants, including *KDM5B, CDC42*, and *CREB5*. SNPs annotated to these miRNA target genes were identified as having the strongest associations with GA (**Figure 4B**). We did not find any significant genetic variants within the primary sequences of these 10 miRNAs.

**Figure 4.**
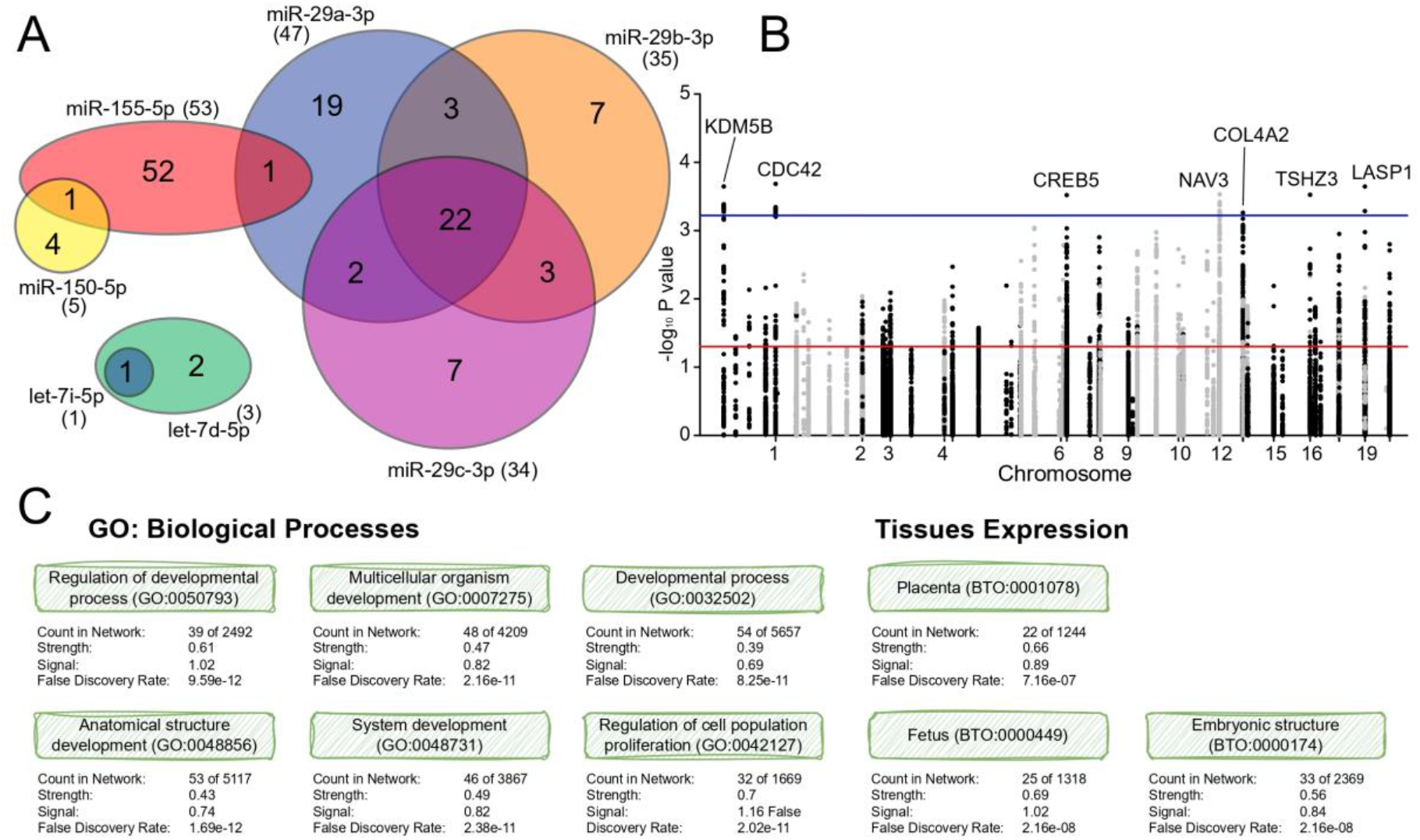
(A) Venn diagram showing overlap of target genes of top 10 miRNAs, identified using miRDB^40^ and validated using miRTarBase^41^; (B) Manhattan plot displaying associations between clinical GA and SNPs in the target genes of top 10 miRNAs, and whether these associations are nominally significant (red line), or are significant after adjusting for multiple testing across the 78 unique target genes (blue line); and (C) gene ontology (GO) terms related to biological processes and tissue expression for the target genes of the top 10 miRNAs, based on STRING^49^. Signal reflects whether the observed number of protein–protein interactions exceeds random expectation, indicating network connectivity. Strength quantifies pathway enrichment as the log-ratio of observed to expected proteins within a functional category, where higher values indicate stronger overrepresentation. An overview of the miRNA target genes and their SNPs is presented in the supplementary materials.

Pathway enrichment analysis for the 78 target genes of the top 10 miRNAs highlighted several significantly associated GO terms and tissue expressions. The strongest associated biological processes included “Regulation of developmental process” and “System development”. Additionally, tissue expression analysis indicated strong associations with fetal, embryonic, and placental tissues, supporting the developmental relevance of these genes (**Figure 4C**). An overview of target genes and SNPs is presented in the supplementary materials.

#### Development of a gestational age miRNA clock (miRClock-GA)

An alpha of 1 consistently resulted in the lowest mean-squared error (MSE) for miRClock-GA. For the training set, the minimal lambda of 0.05 corresponded to a cross-validated MSE of 1.77 (RMSE=1.33), resulting in the inclusion of 53 miRNAs in the model, of which 21 (∼39.6%) were also identified as significant after multiple testing correction in the miRNome-wide association study of Step 1. The correlation between clinical GA and miRClock-GA (**Figure 5**) in the training set was 0.62 (*R*^2^=0.38). For the test set, the model showed a MSE of 1.58 (RMSE=1.26) and a correlation between clinical GA and miRClock-GA of 0.49 (*R*^2^=0.24). The model performance remained consistent over the different train-test splits (Train r_*range*_=0.62 - 0.72; Test r_*range*_=0.45 - 0.52). Sensitivity analyses in complete cases data showed similar results, while there was a slightly higher correlation in the subgroup in which GA was also measured using LMP in both train (r_*US*_=0.62, r_*LMP*_=0.66) and test sets (r_*US*_=0.49, r_*LMP*_=0.54). Benchmark metrics and sensitivity analyses on GA estimation methods are presented in the supplementary materials.

**Figure 5.**
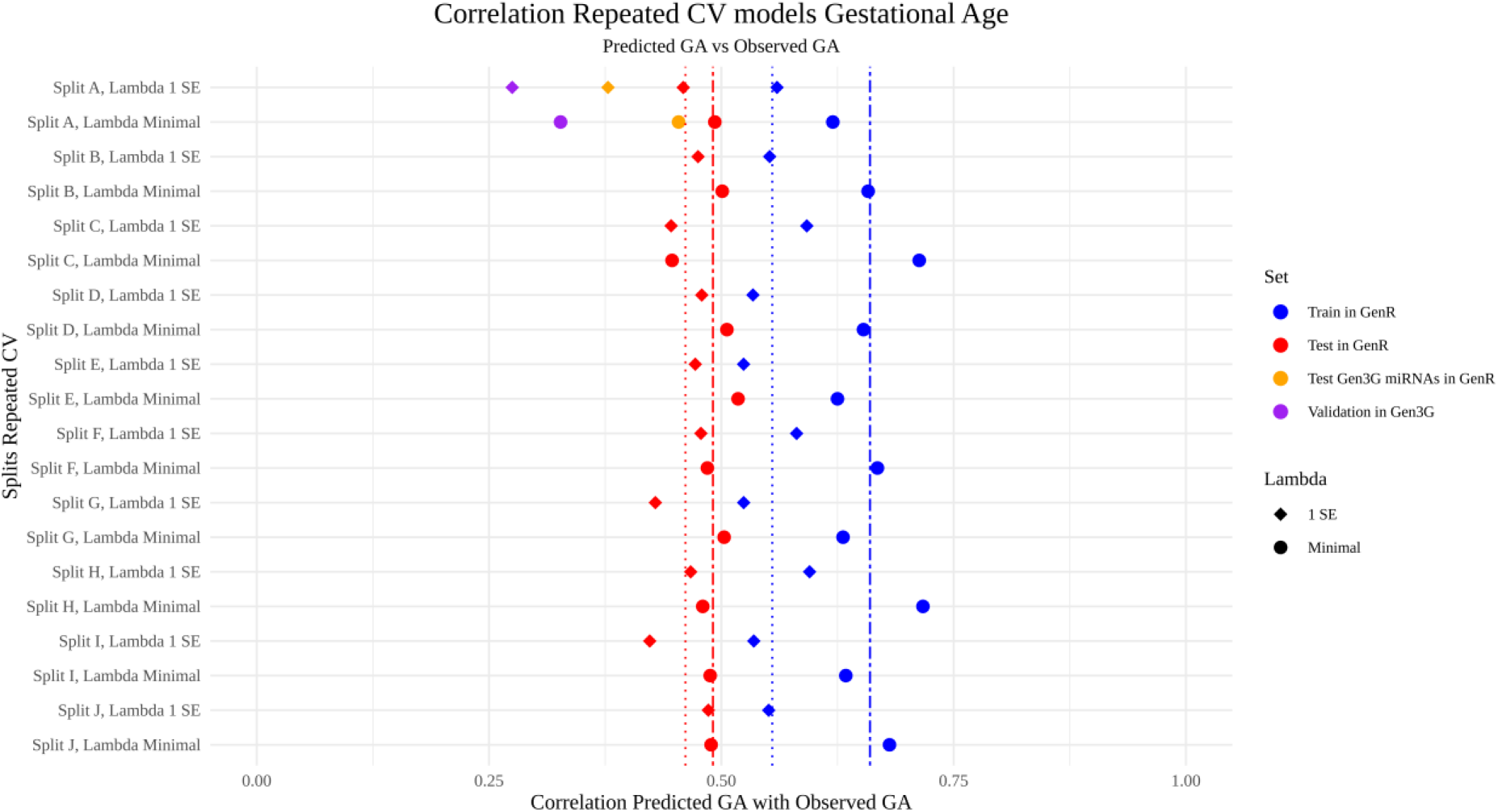
Consistency of correlations between clinical GA and miRClock-GA in the 10x repeated CV model for training and test sets in Generation R; as well as the correlations between clinical GA and miRClock-GA for the test set (Split A) in Generation R based on miRNAs that sufficiently expressed in Gen3G to be included in the calculation of miRClock-GA, and the validation set in Gen3G based on the weights of the first train-test split (Split A). Dashed lines indicate the consistency of correlations between clinical GA and miRClock-GA based on minimal lambda, whereas dotted lines indicate the consistency of correlations between clinical GA and miRClock-GA based on 1 SE lambda. For miRClock-GA in Gen3G, 26 out of 53 miRNAs for the minimal lambda model were available due to minimal expression thresholds, whereas 13 out of 26 miRNAs for the 1 SE lambda model were available.

#### Comparison of gestational age estimates between miRClock and DNAm-based Clocks

In the GenR test set, the pooled Pearson correlation between MiRClock-GA and clinical GA was lower than that observed for DNAmClocks in the same population (r_*miRClock*−*GA*_=0.49, compared to: r_*Bohlin*_=0.73, r_*Knight*_=0.54, and r_*Haftorn*_=0.74; **Figure 6**). Correlations between miRClock-GA and DNAmClock-GA varied from weak to moderate across clocks (r_*Bohlin*_=0.41; r_*Knight*_=0.28; r_*Haftorn*_ =0.42). Correlations between miRClock-AA and DNAmClock-AA were weak(r<.09, P<.05) and not significant with the Knight clock.

**Figure 6.**
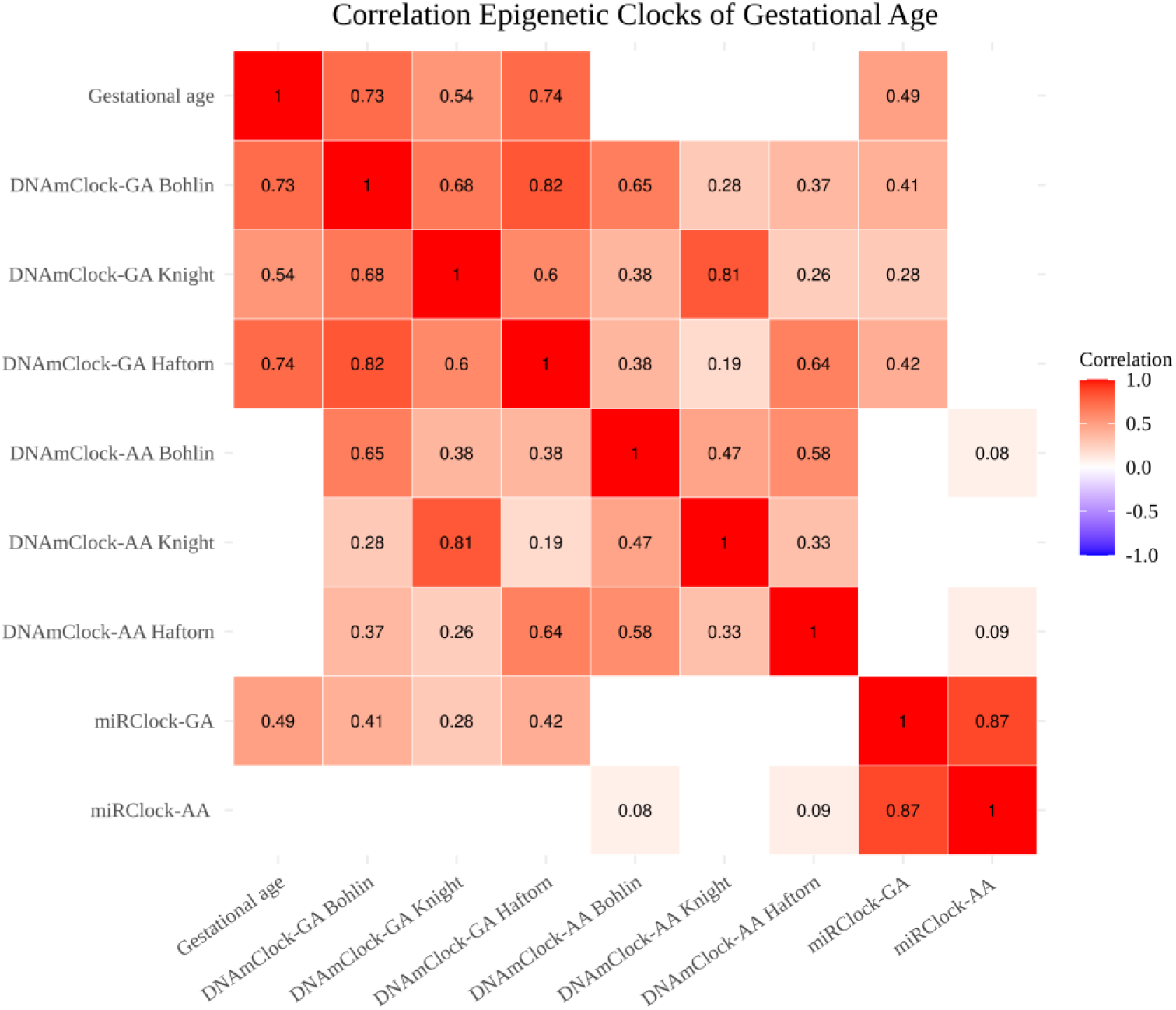
Pooled correlations between clinical GA, miRClock-GA, miRClock-AA, DNAmClock-GA, and DNAmClock-AA in the test set of Generation R (N=678). Correlations not reaching the significance threshold of P<0.05 were left blank.

#### Prospective associations of miRClock with health and developmental outcomes

MiRClock-GA was significantly associated with birthweight after Galwey-correction for multiple testing (P_*cutoff*_=1.32e-03), both in the base model (*R*^2^=0.11, P=4.60e-06) and the extended model, adjusting for a more extensive set of covariates (i.e. maternal smoking, education level, BMI, and age at intake; *R*^2^=0.14, P=1.36e-06). For reference, the model incorporating both covariates and clinical GA explained an *R*^2^ of 0.33 (P=4.96e-46) in the base model and an *R*^2^ of 0.35 (P=7.49e-44) in the extended model. Of DNAmClock-GA, Haftorn performed best in both the base model (*R*^2^=0.22, P=5.30e-24) and the extended model (*R*^2^=0.24, P=3.88e-23), with both Bohlin (Base model: *R*^2^=0.19, P=8.10e-19; Extended model: *R*^2^=0.21, P=8.12e-19) and Knight (Base model: *R*^2^=0.11, P=1.37e-06; Extended model: *R*^2^=0.14, P=3.18e-07) model performance comparable to miRClock-GA. See supplementary materials for further comparisons.

For miRClock-AA, we observed a nominally significant association with birthweight (i.e. independent of clinical GA) in the base model (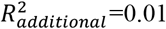, P=3.24e-03) that did not survive multiple testing correction (**Figure 7**), but did in the extended model (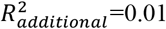, P=1.80e-03). This is likely explained by the inclusion of the maternal covariates in the extended model, which resulted in a reduction in model error, as evidenced by comparable effect sizes yet smaller standard errors. None of the DNAmClock-AA models reached significance after multiple testing correction.

**Figure 7.**
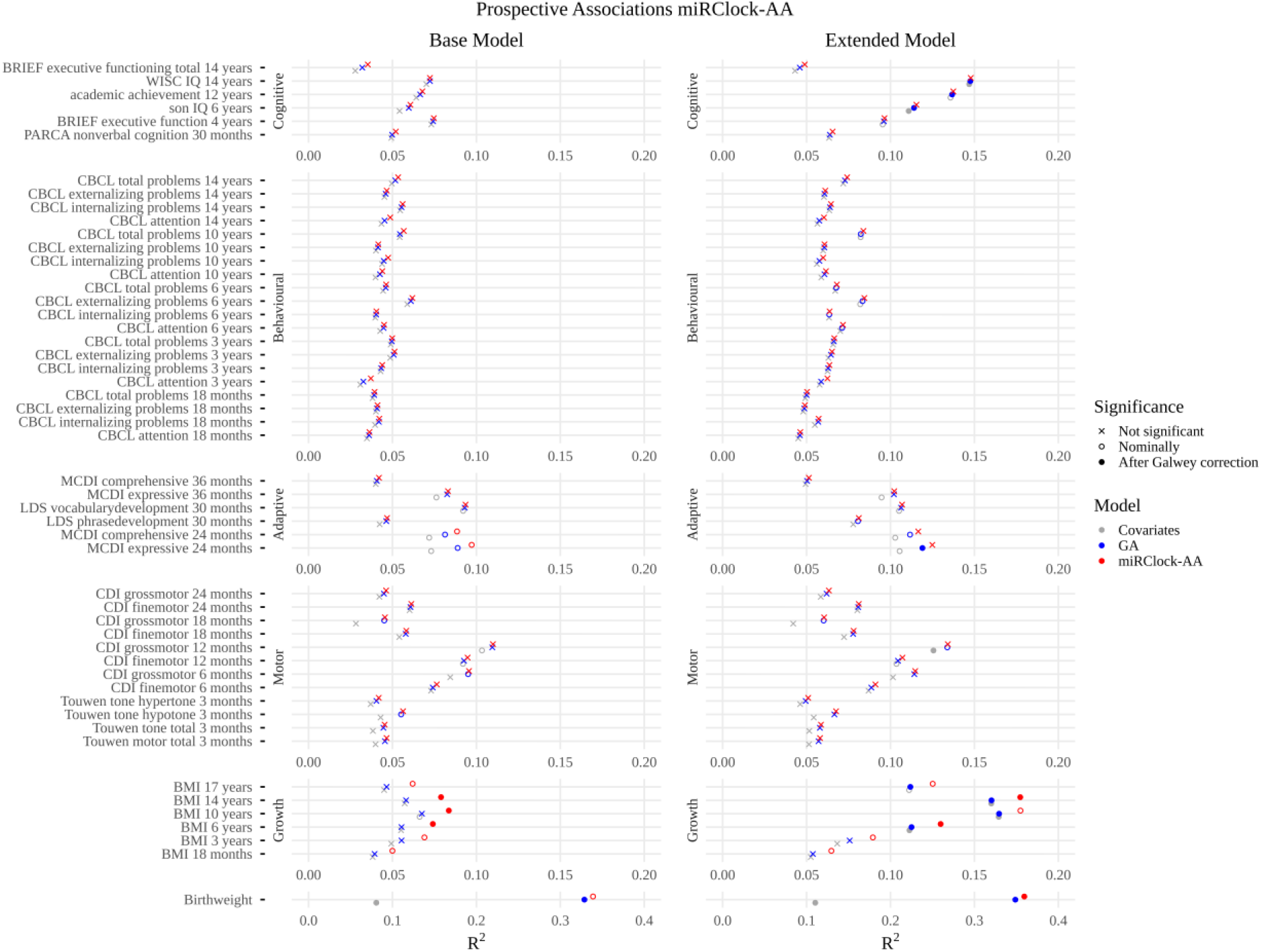
Prospective associations of miRClock-AA with health and developmental phenotypes at later ages. On the left are the base models, on the right the extended models. The extended models included maternal covariates such as maternal age, education level, BMI, and smoking status. The covariate models (in grey) cover cumulative variance explained by sex, haemolysis, miRNA-sequencing plate, initial total sample miRNA concentrations prior to dilution, and blood cell counts as estimated from DNAm data. The GA models (in blue) cover the covariate model in addition to clinical GA. The miRClock-AA models (in red) cover the GA model in addition to miRClock-AA.

Regarding the other developmental outcomes, we found that miRClock-GA was significantly associated with the MacArthur Communicative Development Inventory (MCDI) expressive language subscale at 2y, as well as with BMI at 10y and 14y in the base model, after multiple testing correction. These associations were also observed in the extended models, with the addition of MCDI comprehensive language subscale at 2y, BMI at 6 and 17y, SON IQ at 6y, academic achievement at 12y, and WISC IQ at 14y. For miRClock-AA, we observed three associations in the base model that remained significant after Galwey-correction (**Figure 7**), with BMI at 6 (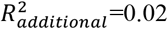, P=1.33e-03), 10 (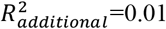, P=3.91e-03) and 14y (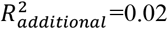, P=8.12e-04). In the extended model, the associations for BMI at 6 (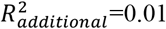, P=1.41e-03) and 14y (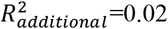, P=1.39e-03) remained significant after Galwey-correction.

#### Validation in the Gen3G cohort

We tested how many of the 123 GA-associated miRNAs could be replicated in an independent cohort of 213 newborns (Gen3G).^25^ Of these, 78 were present in Gen3G with a minimal expression of 5 CPM in at least 50% of samples, and could thus be validated. Effect sizes for the 78 miRNAs between the cohorts were positively correlated (r_*Spearman*_=0.34). 62 miRNAs (79%) were directionally consistent and 13 (17%) showed nominal significance in Gen3G (P<0.05), with one miRNA passing FDR-correction (let-7d-5p, **Figure 3E & F**).

Of the 53 miRNAs included in the calculation of miRClock-GA in Generation R, 26 were present in Gen3G at the minimal expression threshold. As a baseline indication of the influence the limited overlap in miRNAs would have on miRClock-GA (miR^2^_6_Clock-GA), we applied the same set of miRNAs and corresponding weights to the calculation of miR^2^_6_Clock-GA in the Generation R test set, resulting in a MSE of 1.79 (RMSE=1.34). The correlation between clinical GA and miR^2^_6_Clock-GA was 0.45 (*R*^2^=0.21; **Figure 5**), indicating that the missing miRNAs accounted for 3% of the explained variance in miRClock-GA.

Computing miR^2^_6_Clock-GA with weights derived from the GenR training set and applying them to the 26 available miRNAs in Gen3G resulted in a MSE of 1.35 (RMSE=1.16). The correlation between clinical GA and miR^2^_6_Clock-GA was 0.33 (*R*^2^=0.11; **Figure 5**). The improvement in RMSE is likely due to the lack of preterm born children in Gen3G, excluding the samples with the largest deviation between miRClock-GA and GA (**Figure 8**). When we filtered the Generation R test set for samples with a minimal GA of 37 weeks (lowest GA in Gen3G) and tested its performance again, miRClock-GA showed a lower MSE of 1.32 (RMSE=1.15) with a slight drop in its correlation with clinical GA (r=0.44, *R*^2^=0.19).

**Figure 8.**
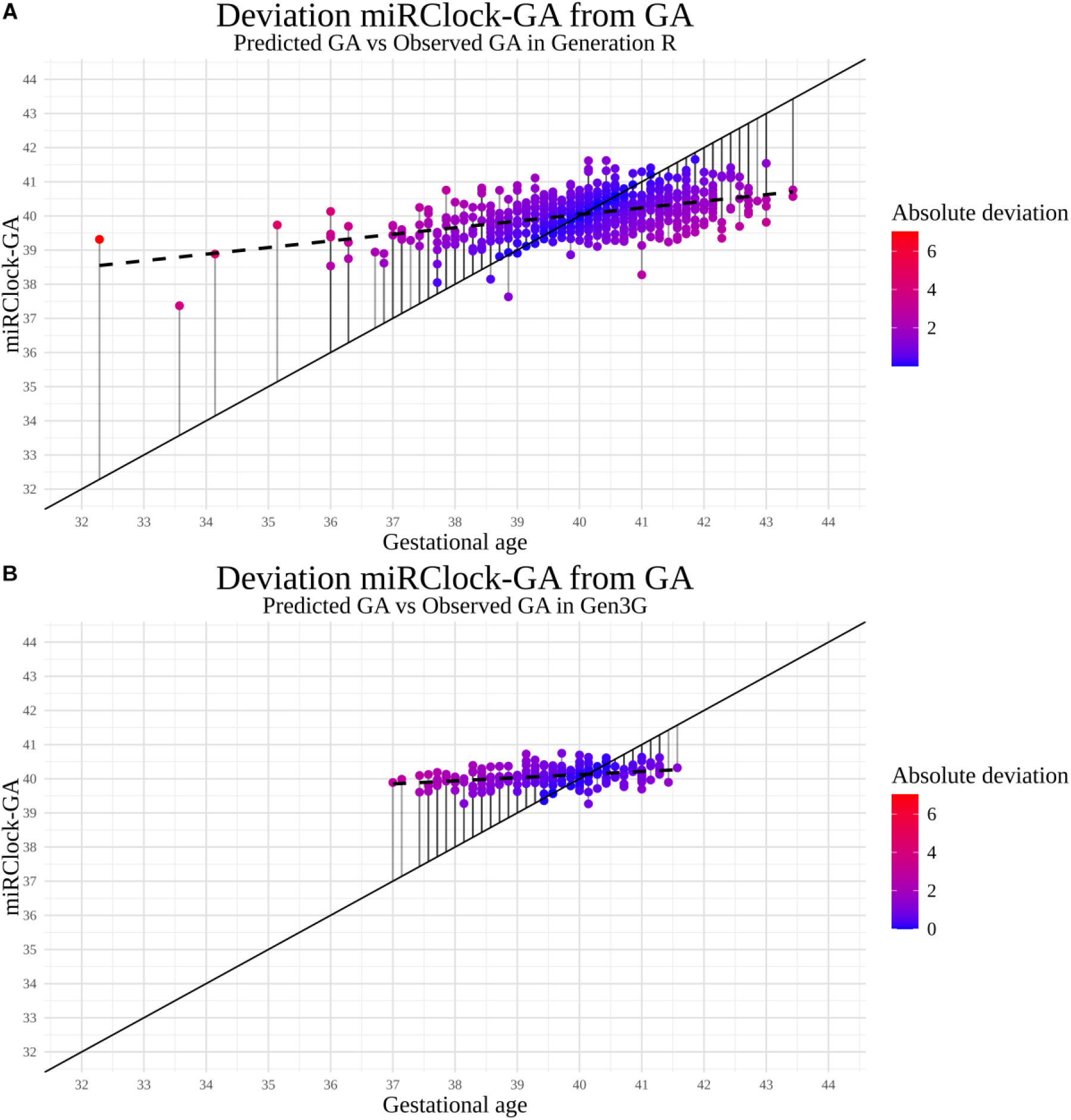
Deviation of miRClock-GA relative to clinical GA for both (A) Generation R- and (B) Gen3G-cohorts. The solid line represents a hypothetical perfect predicted value of GA, with lines from each miRClock-GA value towards this line indicating the corresponding deviation. The colours of the dots indicate the degree of absolute deviation of miRClock-GA from the perfect prediction. The dotted line represents a regression line drawn based on regressing GA onto miRClock-GA.

## Discussion

This is the first study to characterize plasma circulating miRNA profiles associated with gestational age at birth at the population level, using data from 1695 children from Generation R. We highlight here three key findings. First, we found that out of the 2083 miRNAs tested, 123 were significantly associated with gestational age after multiple testing correction. The vast majority (86%) were upregulated with higher gestational age, showing enrichment for developmental processes and fetal tissue expression. Of these GA-associated miRNAs, 78 were also available in an independent cohort of neonates (Gen3G), and tested for validation. Among these 78 miRNAs, 62 (79%) were associated in a directionally consistent manner across cohorts, and 13 (17%) showed nominal significance in Gen3G, with one miRNA passing FDR-correction (let-7d-5p). Second, we created a new epigenetic clock of gestational age – miRClock-GA, which showed moderate correlations with both GA and existing DNAm-based gestational clocks and was validated in Gen3G. Third, we found that miRClock-GA correlated cross-sectionally with birthweight, as well as prospectively with several developmental, growth, and cognitive outcomes measured up to age 17y, after adjusting for a range of maternal covariates. The clock also predicted childhood BMI over and above clinically assessed GA itself.

We found 123 miRNAs significantly associated with gestational age, and none showed significant sex interactions. The top positive association was found for miR-150-5p and the top negative association for miR-373-3p. MiR-150-5p was nominally replicated in Gen3G and has well-established links to pregnancy biology in maternal whole blood as a predictor of premature rupture of membranes-related spontaneous preterm labour through its targeting of chorionic *ADAM19*,^50^ and its impact on placental cellular abilities, e.g. migration, invasion, and angiogenesis of extravillous trophoblast cells.^51^ MiR-373-3p, although not replicated in Gen3G, was found to be more highly expressed in the blood of patients suffering preterm labour, and increasingly so with inflammation, suggesting a role in preterm birth mechanisms.^52^ Additionally, miR-373-3p may play a role in selective intrauterine growth restriction by restricting the growth and migration of HTR8 trophoblast cells through targeting *SLC38A1*.^53^ When validating our findings in Gen3G, 62 (79%) of the 78 available miRNAs were directionally consistent and 13 (17%) miRNAs also showed nominally significant associations with GA in Gen3G, with let-7d-5p surviving multiple testing correction. Upregulation of let-7d-5p has been identified in maternal plasma as early as 12 to 14 weeks of gestation in pregnancies that later resulted in small-for-gestational-age births.^54^ The lack of statistical significance for most tested miRNAs in Gen3G may reflect a number of factors, such as low power - given that Gen3G had a substantially smaller sample size compared to GenR, but effect sizes between cohorts were overall correlated and directionally consistent - as well as sequencing depth differences. Of the 2125 miRNAs measured in Gen3G, 560 both passed the minimal expression threshold (≥5 CPM in ≥50% of samples) and were present in Generation R, restricting the potential overlap to a maximum of 78 eligible miRNAs among the 123 significant findings from Generation R. Interestingly, comparison with miRNAs associated with adult age^19^ revealed no overlap, suggesting that age-related miRNA signatures at birth are largely distinct from those observed in adulthood.

Next, we constructed and validated a miRNA-based GA clock (miRClock-GA), which explained a substantial proportion of the variation in clinical GA and showed consistent performance across train–test splits. A combination of 53 miRNAs was selected to optimally predict GA, almost 40% of which were already identified in the miRNome-wide analysis, suggesting miRClock-GA captures a substantial amount of the variance in GA. In the independent cohort, only 26 of the 53 miRNAs were available in the data at minimal expression thresholds. Yet, in this cohort the restricted clock (miR^2^_6_Clock-GA) also performed moderately well, correlating significantly with GA. Applying this overlapping miRNA subset to the GenR test set to establish a baseline effect of limited miRNA overlap on miRClock-GA, we noted increased predictive precision for term versus preterm births. MiR^2^_6_Clock-GA prediction in Gen3G showed a lower RMSE than miRClock-GA in GenR, even though the correlation with clinical GA was lower in Gen3G compared to GenR. When limiting GenR test set to participants with a minimal GA of 37 weeks, as present in Gen3G, this inconsistency dissolved. This finding suggests miRClock-GA shows better prediction for term births than for preterm births.

When relating miRClock-GA to three commonly used DNAm-based clocks, we found that all clocks were significantly intercorrelated, although the magnitude of correlations was smaller across modalities (miRNA and DNAm) than within modality (i.e. among DNAm-based clocks). This suggests miRClock-GA may capture partially distinct, complementary, biological signals, that potentially reflect different exposures, epigenetic- or developmental processes relative to DNAmClocks. We also found that DNAm-based clocks showed stronger correlations with clinical GA as compared to miRClock-GA. These differences could stem from biological (e.g. relative role of miRNA vs DNAm-based regulation in GA) or technical factors (e.g. model training cohort, number of features using DNAm data vs miRNA data, stability of GA-relevant signatures, sequencing depth, normalization approach, or residual batch effects) affecting the estimated GA. Future comparisons between miRClock-GA and DNAmClock-GA may be directed at tissue or cell-type specificity to explore whether miRNAs used for calculating miRClock-GA are enriched in specific fetal or placental cell types, relative to CpGs driving DNAmClocks. Additionally, testing the strength of miRClock-GA and DNAmClock-GA, and corresponding miRNAs and CpGs, in associations with prenatal exposures, e.g. maternal smoking, inflammation, BMI, stress, may indicate the degree to which these clocks are responsive to prenatal and intrauterine environmental influences. Alternatively, individuals with large discrepancies between miRClock-GA and DNAmClock-GA could be tested on whether discordant profiles associate with specific exposures or phenotypes.

Finally, we examined how miRClock-GA related to developmental outcomes from birth to adolescence. As expected, miRClock-GA correlated cross-sectionally with birthweight, a phenotype strongly associated with clinical GA. Additionally, miRClock-GA associated with markers of growth, adaptive, and cognitive development. All epigenetic clocks displayed prospective associations largely consistent with those of clinical GA, supporting their capacity to capture shared developmental variance. When we examined whether miRClock-GA explains any variance beyond clinical GA itself (miRClock-AA), we found that miRClock-AA correlated with birthweight and childhood BMI measures. Inclusion of maternal covariates in the extended model resulted in a reduction of the model error, making the prediction of the model more precise. These results are in line with previous findings relating epigenetic gestational age acceleration to trajectories of height and weight during childhood.^55^

Contrary to miRClock-AA, none of the three DNAmClock-AA showed significant associations with prospective outcomes in GenR data. Consistent with Salontaji et al., who reported that DNAmClock-AA provided little additional information about ADHD risk beyond clinical GA,^13^ our findings here similarly suggest limited added predictive value of accelerated ageing. The absence of significant prospective associations, with the exception of miRClock-AA with childhood BMI, suggests limited utility for capturing more than clinical GA itself, i.e. acceleration/deceleration, and related downstream phenotypes. Adult studies have shown how second (i.e. trained on age-related phenotypes, mortality) and third (i.e. trained on ageing trajectories) generation DNAmClocks outperform first generation DNAmClocks (i.e. trained on chronological age) in predicting age-related outcomes.^56^ We may see similar developments for miRNA-based GA clocks as well, where clocks are trained on other measures than chronological age (e.g. biological maturation, longitudinal pre- or postnatal growth measures) to estimate prospective health- and developmental outcomes.

Strengths of this study include leveraging to our knowledge the largest population birth sample miRNA dataset to characterize miRNAs associated with GA, to produce a miRNA-based GA epigenetic clock, validate findings in an independent sample, and incorporate a wide set of longitudinal developmental outcomes spanning birth to adolescence. However, several limitations should be considered. First, participants included in the miRNA analyses were selected based on continued participation across data collection waves, which may have introduced “healthy volunteer”-selection bias towards healthier and more socioeconomically advantaged individuals.^57^ This may have potentially reduced variability in GA and developmental outcomes (**Supplementary Figure S10**), thus reducing power to find prospective associations with both GA and miRClock-AA in our sample relative to those in the general population, and underestimating any found associations (e.g. associations could be stronger in children with higher risk profiles). Second, the study population was restricted to individuals of European ancestry, necessitating future studies to test generalizability to other ancestral backgrounds. Finally, although this represents the largest study of its kind to date, the modest effect sizes inherent to circulating miRNAs suggest that we may still have been underpowered to detect more subtle associations, consistent with the observation that several clock-selected miRNAs did not survive multiple-testing correction in miRNome-wide analyses.

In conclusion, we identified 123 miRNAs significantly associated with gestational age, with our top miRNAs linked to fetal tissues and prenatal health. Associations were directionally consistent and partially validated in an independent birth sample, and largely distinct from miRNAs associated with chronological age in adulthood. Using the miRNA data, we built an epigenetic clock of GA (miRClock-GA) explaining a substantial proportion of the variation in clinical GA. This clock was independently validated, and captures signal that appears to be partly distinct from established DNAm-based clocks (Bohlin, Knight, and Haftorn). The clock also explained variance in birthweight and BMI across development beyond clinical estimates of GA (i.e. miRClock-AA) after adjustment for developmental covariates such as maternal age, education level, smoking, and BMI. This may point to applicability of miRClock-AA to outcomes also assessed at the time of birth and highlights circulating miRNAs as potential markers of gestational biological ageing complementary to DNAm-based markers.

## Supporting information

Manuscript Supplement

## Acknowledgements

The Generation R Study is conducted by Erasmus MC, University Medical Center Rotterdam in close collaboration with the School of Law and Faculty of Social Sciences of the Erasmus University Rotterdam, the Municipal Health Service Rotterdam area, Rotterdam, the Rotterdam Homecare Foundation, Rotterdam and the Stichting Trombosedienst & Artsenlaboratorium Rijnmond (STAR-MDC), Rotterdam. We gratefully acknowledge the contribution of children and parents, general practitioners, hospitals, midwives and pharmacies in Rotterdam. The study protocol was approved by the Medical Ethical Committee of Erasmus MC, Rotterdam. Written informed consent was obtained for all participants. We gratefully acknowledge the contribution of the team performing the microRNA sequencing using the HTG EdgeSeq miRNA Whole Transcriptome Assay for the Generation R Study, as executed by the Human Genotyping Facility of the Genetic Laboratory of the department of Internal Medicine, Erasmus MC, the Netherlands. We thank Mr. Pascal Arp, Mr. Ramazan Buyukcelik, Mr. Vid Prijatelj, and Ms. Anna Ku.

The general design of the Generation R Study is made possible by financial support from Erasmus MC, Erasmus University Rotterdam, the Netherlands Organization for Health Research and Development and the Ministry of Health, Welfare and Sport. The work of TF, AN, and CAMC was supported the European Union’s Horizon Europe Research and Innovation Programme (FAMILY, No.101057529). AN and CAMC are supported by the European Research Council (TEMPO, No.101039672). This research was conducted while CAMC was a Hevolution/AFAR New Investigator Awardee in Aging Biology and Geroscience Research.

This project has received funding from the European Union’s Horizon Europe Research and Innovation Programme (STAGE, No.101137146). Views and opinions expressed are however those of the author(s) only and do not necessarily reflect those of the European Union. Neither the European Union nor the granting authority can be held responsible for them.

The work of MG was partly supported by the Erasmus MC Incentive grant (No.118110) and Alzheimer Nederland grant (WE.06-2025-04). The mentioned funders had no role in the design and conduct of the study, nor in the decision to submit the manuscript for publication.

Gen3G was initially funded by the Fonds de recherche du Québec (FRQ)—Santé operating grant (to M-FH, grant #20697); a Canadian Institute of Health Research (CIHR) operating grants (to M-FH grant #MOP 115071 and to LB grants #PJT152989, IGH155183 and PJT190076), Diabète Québec, Internal funding supports from le Centre de recherche du CHUS and l’Université de Sherbrooke. The measurements of cord blood microRNA was supported by the FRQ-Réseau de recherche en santé cardiométabolique, diabète et obésité.

Views and opinions expressed are however those of the author(s) only and do not necessarily reflect those of the European Union or the European Health and Digital Executive Agency. Neither the European Union nor the granting authority can be held responsible for them.

## Data sharing statement

Full summary statistics are included in the supplementary materials. The preregistration of the study protocol can be found on the website of Open Science Foundation (https://osf.io/7yb4m/overview). The data that support the findings of this study are not publicly available due to privacy and legal restrictions but are available upon reasonable request to the head of the Generation R Study: and https://www.generationr.nl, subject to local, national, and European rules and regulations.

In Gen3G, pregnancy miRNA seq data are on GEO, accession #GSE216998. Currently we are in the process of adding the cord blood miRNA data to this account.

## Declaration of interests

The authors declared no competing interests.

## Contributors

Conceptualisation: TF, AN, CAMC

Data curation: TF, MMK, MG, AN, CAMC,

JFF Formal analysis GenR: TF, MMK

Formal analysis Gen3G: FW

Funding acquisition: NvH, AN, CAMC, MG, LB

Methodology: TF, AN, MG, CAMC

Project administration: CAMC

Supervision: AN, MG, CAMC, MFH, PEJ

Writing - original draft: TF, MMK, FW, MG, AN, CAMC

Writing - review & editing: All authors

Data interpretation: all authors

All authors read and approved the final version of the manuscript.

## Notes

### Competing Interest Statement

The authors have declared no competing interest.

### Summary of Updates

Original PDF version of the manuscript had problems with in-text symbols for beta, r, and R^2. This was fixed for the new version.

