## Supplementary material for "Development of a microRNA clock for gestational age in the general population": Manuscript Supplement

Development of a microRNA clock for gestational age in the general population. Supplementary methods and results.

a. Department of Child and Adolescent Psychiatry, Erasmus MC, University Medical Center Rotterdam, Dr. Molewaterplein 40, 3015 GD Rotterdam, the Netherlands
b. Generation R Study Group, Erasmus MC, University Medical Center Rotterdam, Dr. Molewaterplein 40, 3015 GD Rotterdam, the Netherlands
c. Department of Epidemiology, Erasmus MC, University Medical Center Rotterdam, Dr. Molewaterplein 40, 3015 GD Rotterdam, the Netherlands
a. Département de Biologie, Université de Sherbrooke, Sherbrooke, QC, Canada
d. Department of Pediatrics, Erasmus MC, University Medical Center Rotterdam, Dr. Molewaterplein 40, 3015 GD Rotterdam, the Netherlands
a. Genomics Core Facility, Erasmus MC, University Medical Center Rotterdam, 3015 Rotterdam, The Netherlands
e. Department of Social and Behavioral Sciences, Harvard T. H. Chan School of Public Health, Harvard University, Boston, Massachusetts, USA
a. Centre de recherche et d'innovation, Centre Intégré Universitaire de Santé et de Services Sociaux (CIUSSS) du Saguenay-Lac-Saint-Jean, Saguenay, Québec, Canada
b. Département Clinique de Médecine de Laboratoire, Centre Intégré Universitaire de Santé et de Services Sociaux (CIUSSS) du Saguenay-Lac-Saint-Jean, Hôpital de Chicoutimi, Saguenay, Québec, Canada
c. Université du Québec à Chicoutimi, Saguenay, Québec, Canada
a. Centre de Recherche du Centre Hospitalier Universitaire de Sherbrooke (CRCHUS), Sherbrooke, QC, J1H 5N3, Canada
b. Centre de recherche du Centre hospitalier universitaire de Sherbrooke (CRCHUS), Sherbrooke, QC, Canada
f. Division of Chronic Disease Research Across the Lifecourse (CoRAL), Department of Population Medicine, Harvard Medical School, Harvard Pilgrim Health Care Institute, Boston, MA, USA
g. Diabetes Unit, Massachusetts General Hospital, Boston, MA, USA

* Shared first author, # Shared senior author

### Supplementary methods

#### Measures in Generation R

**MiRNAs expression profiling and normalization in Generation R**. Plasma circulating miRNA was collected from umbilical cord blood at birth and frozen at -80 °C. The samples of 1710 participants were randomized across 18 plates, consisting of 95 participant samples and one control sample on each plate. Samples were thawed and subsequently centrifuged and separated for next-generation sequence analysis through use of HTG EdgeSeq miRNA Whole Transcriptome Assay in accordance with the manufacturer’s instructions. These are described in more detail elsewhere.^1^ In short, 15 µL of plasma was mixed with an equal volume of lysis buffer, followed by the addition of Proteinase K (1/10th of lysate volume) and incubation at 50°C for 180 minutes. Prepared samples (25 µL per well) were pipetted into the HTG barcoded sample plate, with probe capture running for 20 hours. Samples were then stored at −20°C or processed immediately. For PCR, a 21 µL master mix was used, with cycling conditions: 95°C for 4 min, 16 cycles of 95°C for 15 sec, 56°C for 45 sec, and 68°C for 45 sec, followed by 68°C for 10 min and hold at 4°C. Library cleanup involved incubating 37.5 μL of AMPure XP beads (Beckman Coulter, Inc., Pasadena, California, USA) with 15 μL of PCR-amplified libraries. Libraries were quantified using the KAPA Library Quant kit (Roche, Basel, Switzerland) and qPCR (95°C for 5 min, 35 cycles of 95°C for 30 sec and 60°C for 45 sec). The HTG Library Calculator (v1.3) determined final dilution volumes. Diluted libraries were pooled, spiked with PhiX control libraries, and loaded into the NextSeq 2000 P2 flow cell.

Sequencing was performed on NextSeq 2000 (Illumina Inc., San Diego, California, USA) using a single end 50 base-pair approach. Demultiplexing was done with bcl-convert in DRAGEN Host Software (Version 07.021.624.3.10.11), and data quality and integrity was assessed using [Illumina Sequencing Analysis Viewer, with >90% of bases exceeding PHRED Q30](https://emea.illumina.com/systems).^2^ HTG EdgeSeq parser software processed fastq files, aligning reads to a custom FASTA file containing 2102 sequences. Bowtie2 aligner was used to ensure a non-greedy, conservative alignment,^3^ with an overall alignment rate >85%, independently replicated in our compute environment. The miRNA probe list was based on miRBase v20.^4^
Initial post-sequencing quality control (QC) of aligned data was performed using HTG EdgeSeq spreadsheet developed for Microsoft Excel (Microsoft Inc., Redmond, Washington, USA). Three metrics were checked: RNA degradation or poor quality (QC0), low read depth (QC1), and non-biological library variability (QC2). While no samples failed QC0, one participant sample and one control sample failed QC1 and 14 participant samples as well as 2 control samples failed QC2. This resulted in a final dataset of 1695 participant samples with expression data for 2083 miRNAs.

Read counts were first adjusted for total reads within each sample, after which they were normalized using trimmed mean of M-values (TMM),^5^ scaled to counts per million (CPM), and subsequently log-transformed. In order to mitigate vulnerabilities of TMM-normalized RNAseq counts to exaggerated false positive findings^6^, we applied 95% winsorization as recommended by^7^.

**Developmental outcomes from birth to adolescence.** To assess the association of miRClock-GA with later child development we focused on 51 outcomes categorized along 5 developmental domains, encompassing (i) growth, (ii) motor, (iii) adaptive, (iv) behavioural, and (v) cognitive development.

i. **Growth developmental outcomes**
   *Infant Growth.* Length, height and weight were measured at birth, and at 1.5, 3, 6, 10, 14, and 17 years old. From length and weight, BMI was calculated and transformed into age- and sex-adjusted SD-scores.
ii. **Motor developmental outcomes**
    *Touwen’s Neurodevelopmental Examination (TNE).* Infant motor development was administered by trained research assistants during a home visit at 3 months of age using a modified version of TNE.^8^ The TNE was assessed across across four scales; motor total, tone total, hypotone, and hypertone. Active and passive muscle tone assessments were added, following the method of de Groot.^9^ Tone was assessed in supine, horizontal, vertical, prone, and sitting positions, with items scored as low, normal, or high. Response behaviour was observed in supine, prone, sitting, standing, and walking positions, with items scored as absent, present, or excessive. Other observations were scored as absent, present, or excessive. Overall, age-normative responses were labelled as “optimal”, whereas responses were labelled as “nonoptimal” when the response indicated a delayed development (optimal = 0, non-optimal = 1). Scale values were calculated by summing non-optimal items, with a lower number indicating a more appropriate neurodevelopment.
    *Child Development Inventory (CDI) - Gross and Fine Motor Development.* The CDI was main-caregiver reported at 6 months as well as at 1, 1.5, and 2 years. The CDI assesses fine and gross motor development.^10^ The Fine Motor Development scale measures eye-hand coordination. The scale contains dichotomous yes/no items (e.g. “Builds a tower of two or more blocks”, “Scribbles with crayon or pencil”). The Gross Motor Development scale measures walking, running, climbing, jumping, riding, balance, and coordination. Total fine and gross motor development raw scores were standardized using normative data derived from the CDI manual.
iii. **Adaptive developmental outcomes**
     *MacArthur Communicative Development Inventory (MCDI).* The MCDI was reported on by main caregivers at 1.5 years old. Receptive and expressive language was assessed using 112 words selected from the original MCDI.^11^ Main caregivers reported on two sections:
1. Receptive language: yes/no-items to assessing the child’s understanding of short sentences, e.g., “Give me a kiss” and “Say hello”.
2. Expressive language: the child’s production and comprehension of monomorphic root words, rated on a 2-point Likert scale (1 = understands, 2 = says), e.g., “Vroom vroom (car)” and “Camera”.
   Per the scoring protocol, if three consecutive items were marked “no,” all subsequent items were coded as “no” as well. A total score was calculated by summing yes-responses.
   *Language Development Survey (LDS).* The LDS was reported on by main caregivers at 2.5 years old. Receptive and expressive language was assessed in a Dutch translation of the LDS, consisting of 310 vocabulary items (e.g., “Apple” and “Tortoise”) divided into 14 semantic categories.^12^ For each word, caregivers indicated whether the child used it spontaneously (yes/no). Standardized scores were derived from raw scores using the normative data provided in the LDS manual.
iv. **Behavioural developmental outcomes**
    *Child Behaviour Checklist (CBCL).* The CBCL was administered by parents at 1.5, 3, 6, 10, and 14 years of age to assess a child’s behavioural and emotional functioning. Parents rated items on the checklist on a 3-point scale: 0 (“not true”), 1 (“somewhat or sometimes true”), and 2 (“very true or often true”). The responses are scored to generate raw scores for a total score as well as subscales relating to attention problems, externalizing problems, and internalizing problems.
v. **Cognitive developmental outcomes**
   *Total academic achievement.* School performance at 12 years old is measured with a test at the end of primary school created by the Central Institute for Test Development (CITO).^13^ Test scores ranging from 501 to 550 were transformed to percentile rank scores.
   *Non-verbal cognitive development.* Non-verbal cognitive development was reported by main caregiver when the child was 2.5 years old, using the Dutch version of the Parent Report of Children’s Abilities (PARCA).^14^ The PARCA assesses quantitative skills, spatial abilities, symbolic play, planning and organizing, adaptive behaviors, and memory using 26 yes/no items (0 = no, 1 = yes), for example “Can your child put a simple piece, such as a square or an animal, into the correct piece on a puzzle board?”. Items are summed to provide a total score for non-verbal cognitive development.^15^ Raw scores were standardized using normative data provided in the PARCA manual.
   *Executive functioning.* Executive functioning was reported by main caregiver at 4 and 14 years of age using the Brief Rating Inventory of Executive Function-Preschool Version (BRIEF-P).^16^ The BRIEF-P contains 63 items from which a total executive composite score is derived, where higher scores reflect more difficulties in executive functioning. Each item is rated on a 3-point Likert scale (1 = never, 2 = sometimes, 3 = often). Example items include “Has difficulty putting a brake on his/her behaviour, even if asked to do that”, “Can not take changes in routine, eating, places etc.” and “Reacts more intensely to situations than other children”. Standardized scores are derived from the raw data using normative information provided in the manual.
   *Non-verbal intelligence.* Non-verbal intelligence was assessed by trained examiners at 6 years of age, using two subtests from the Dutch non-verbal intelligence test, Snijders-Oomen Niet-verbale intelligentie test 2.5-7 jaar Revisie (SON-R 2.5-7).^17^ Raw scores were summed and adjusted to produce age-normed Total IQ scores.
   *Intelligence.* Intelligence was assessed by trained examiners at 14 years of age, using an abbreviated version of the Wechsler Intelligence Scale for Children (WISC-V),^18^ which is a widely used psychological assessment tool designed to measure intellectual ability of children aged 6 to 16 years. Items were summed into raw scores, from which an age-normed IQ score was derived.

**DNA methylation data**. The processing, control steps, and normalization procedures of DNA methylation data in Generation R is described extensively elsewhere.^19–22^ In short, Generation R umbilical cord blood samples were processed using either the HumanMethylation450K BeadChip array or the MethylationEPIC v1.0 BeadChip. The CPACOR workflow was used for quality control and normalization,^23^ during which arrays with observed technical problems were removed (e.g. failed bisulfite conversion, hybridization, extension problems, sex mismatch as determined by X and Y chromosome probe intensities). Furthermore, probes with a detection P-value above background (P>1.0e-16) were set to missing. Arrays with a call rate >95% per sample were included and quantile normalized.

We examined three DNAm-based GA clocks that are commonly used and have been previously calculated in Generation R.^21^ These include the Bohlin clock (trained on Illumina 450K array in MoBa),^24^ the Knight clock (trained on Illumina 27K and 450K arrays from six cohorts),^25^ and the Haftorn clock (trained on EPIC 850K array in MoBa).^26^ For each clock, we calculated epigenetic age acceleration, which reflects the residual after adjusting epigenetic age for clinical GA.

Blood cell counts were estimated using the combined cord blood-specific reference set in the “FlowSorted.CordBlood.Combined.450K”-Bioconductor package.^27^ This resulted in DNAm-based blood cell proportions for B cells, CD4+ T cells, CD8+ T cells, granulocytes, monocytes, natural killer cells, and nucleated red blood cells. For the analyses featuring DNAm-based clocks, we also included methylation array plate as a measure of batch effects in the DNAm data.

#### Measures in Gen3G

**MiRNAs expression profiling and normalization**.

Total RNA was extracted from 130–500 $\mu$L of cord blood plasma (depending on sample availability) using the mirVana PARIS Kit (ThermoFisher Scientific, catalog #AM1556) according to the manufacturer’s protocol. Prior to extraction, samples were randomized to minimize batch effects, and haemolysis status was assessed using a visual analogue scale ranging from 0 (non-haemolytic) to 3 (highly haemolytic). Total RNA was eluted in 75 µL of nuclease-free water. Prior to library preparation, 70 µL of RNA eluate was concentrated by ethanol precipitation as previously described.^28^ Briefly, 35 µL of cold (4°C) 7 M ammonium acetate solution (ThermoFisher Scientific, catalog #02002268) was added to the RNA sample, followed by 420 µL of pre-chilled (−20°C) absolute ethanol (Commercial Alcohols, ON, Canada; catalog #P006EAAN). The mixture was incubated overnight at −20°C to precipitate RNA, then centrifuged at 16000 × g at 4°C for 30 minutes. The resulting RNA pellet was washed twice with 200 µL of 80% ethanol, centrifuging at 16000 × g at 4°C for 5 minutes after each wash. The pellet was air-dried at room temperature for 30 minutes and resuspended in 5 µL of nuclease-free water. The entire 5 µL volume was used for library preparation. Sequencing libraries were prepared using the TruSeq Small RNA Sample Prep Kit (Illumina, BC, Canada, catalog #RS-200-0012) following the low-input RNA protocol described by Burgos et al.^28^

After randomization of samples, the protocol involved sequential ligation of 3′ and 5′ adapters to small RNAs, reverse transcription to generate cDNA, and PCR amplification (15 cycles) with indexed primers. To optimize the reaction ratio for low RNA concentrations, reagent volumes were reduced to half of the manufacturer’s recommended amounts. Each sample was tagged with a unique index (indexes 1–48, one per sample) to enable multiplexed sequencing. Following amplification, libraries were size-selected by gel electrophoresis on 6% Novex polyacrylamide TBE gels (ThermoFisher Scientific, catalog #EC265BOX). Products in the 145–160 bp size range (corresponding to microRNAs plus adapter sequences) were excised from the gel and purified by overnight elution in 300 µL of nuclease-free water at room temperature with agitation (500 RPM) on a microplate shaker (VWR, catalog #12620-930). Eluted libraries were concentrated by ethanol precipitation following the manufacturer’s instructions, including a 30-minute incubation at −80°C, and resuspended in 25 µL of 10 mM Tris-HCl (pH 8.5). Library quantification was performed using the KAPA Library Quantification Kit for Illumina platforms (KAPA Biosystems, Wilmington, United States) with revised primers and SYBR Fast Universal kit. Libraries were normalized and pooled equimolarly in batches of 48 libraries per lane at a final concentration of 225 pM. Pooled libraries were denatured and clustered on an Illumina NovaSeq S1 flow cell following the Xp protocol according to the manufacturer’s recommendations. Single-end sequencing was performed for 100 cycles on the NovaSeq 6000 platform at the Centre d’expertise et de services Génome Québec (Montréal, Canada).

Raw FASTQ files were processed using the exceRpt small RNA-seq pipeline with minor adaptations.^29^ Rather than the default fastx_clipper, adapter sequences were trimmed using Cutadapt^30^ to enable filtering by maximum read length, which was set to 30 nucleotides. Reads mapping to potential contaminants, including UniVec sequences and ribosomal RNA, were removed. Remaining reads were aligned to the human reference genome (GRCh38) and transcriptome using STAR aligner.^31^ MiRNA quantification was performed by the exceRpt pipeline, which assigns reads to miRNA annotations mainly based on miRBase (release 21).

From the initial cohort, we retained samples with both DNAm and small RNA sequencing data available from cord blood. Siblings were excluded to ensure sample independence. Principal component analysis was performed on the miRNA expression matrix to identify outliers. As a results, three samples with aberrantly low numbers of detected miRNAs were identified as outliers and excluded from subsequent analyses. This resulted in a dataset consisting of 213 samples with expression data for 2125 miRNAs. Due to the large number lowly expressed miRNAs, miRNAs with a minimum of 5 CPM in at least 50% of participants were retained. After filtering, the 680 remaining miRNAs were normalized following the same procedure as in the Generation R Study.

#### Software Generation R

The scripts used for data cleaning and QC in Generation R [(https://github.com/TimFinke/miRNA_QC)](https://github.com/TimFinke/miRNA_QC) as well as those used for the analyses can be found on GitHub [(https://github.com/TimFinke/miRNA_clock)](https://github.com/TimFinke/miRNA_clock). During our analysis we used R^Version 4.5.2; 32^ and the R-packages *bookdown*^Version 0.46; 33^, *broom*^Version 1.0.12; 34^, *dplyr*^Version 1.2.0; 35^, *forcats*^Version 1.0.1; 36^, *ggh4x*^Version 0.3.1; 37^, *ggplot2*^Version 4.0.2; 38^, *ggrepel*^Version 0.9.6; 39^, *glmnet*^40,41^, *glmnetUtils*^41^, *haven*^Version 2.5.5; 42^, *lubridate*^Version 1.9.5; 43^, *Matrix*^Version 1.7.4; 44^, *Metrics*^45^, *mice*^46^, *mitools*^47^, *openxlsx*^Version 4.2.8.1; 48^, *patchwork*^Version 1.3.2; 49^, *poolr*^50^, *psych*^Version 2.6.1; 51^, *purrr*^Version 1.2.1; 52^, *readr*^Version 2.2.0; 53^, *stringr*^Version 1.6.0; 54^, *tibble*^Version 3.3.1; 55^, *tidyr*^Version 1.3.2; 56^, *tidyverse*^Version 2.0.0; 57^, *viridis*^58,59^, and *viridisLite*^Version 0.4.3; 59^.

#### Software in Gen3G

The scripts used for data cleaning and QC in Gen3G [(https://github.com/TimFinke/miRNA_QC)](https://github.com/TimFinke/miRNA_QC) as well as those used for the analyses can be found on GitHub [(https://github.com/whitef19)](https://github.com/whitef19). During our analysis we used R^Version 4.5.2; 32^ and the R-packages *bookdown*^Version 0.46; 33^, *broom*^Version 1.0.12; 34^, *dplyr*^Version 1.2.0; 35^, *forcats*^Version 1.0.1; 36^, *ggplot2*^Version 4.0.2; 38^, *ggrepel*^Version 0.9.6; 39^, *glmnet*^40,41^, *glmnetUtils*^41^, *haven*^Version 2.5.5; 42^, *lubridate*^Version 1.9.5; 43^, *Matrix*^Version 1.7.4; 44^, *mice*^46^, *mitools*^47^, *openxlsx*^Version 4.2.8.1; 48^, *patchwork*^Version 1.3.2; 49^, *poolr*^50^, *psych*^Version 2.6.1; 51^, *purrr*^Version 1.2.1; 52^, *readr*^Version 2.2.0; 53^, *stringr*^Version 1.6.0; 54^, *tibble*^Version 3.3.1; 55^, *tidyr*^Version 1.3.2; 56^, and *tidyverse*^Version 2.0.0; 57^.

### Supplementary results

##### MiRClock-GA based on 1se lambda

An alpha of 1 consistently resulted in the lowest mean-squared error (MSE) for miRClock-GA. For the training set, the 1se lambda of 0.09 corresponded to a MSE of 1.89 (RMSE=1.38), resulting in the inclusion of 26 miRNAs in the model, of which 16 (~61.5%) were also identified as significant after multiple testing correction in the miRNome-wide association study of Step 1. The correlation between chronological GA and miRClock-GA in the training set was 0.56 ($R^{2}$=0.31). For the test set, the model showed a MSE of 1.7 (RMSE=1.3) and a correlation between chronological GA and miRClock-GA of 0.46 ($R^{2}$=0.21). The model performance remained consistent over the different train-test splits, as indicated by consistent associations between miRClock-GA in train- (r${}_{range}$=0.52 - 0.6) and test sets (r${}_{range}$=0.42 - 0.49). Sensitivity analyses showed a slightly higher correlation when GA was measured using LMP (in the subsample with high-confidence estimation) as opposed to ultrasound in both train (r${}_{US}$=0.56, r${}_{LMP}$=0.62) and test sets (r${}_{US}$=0.46, r${}_{LMP}$=0.51).

Applying the GenR trained weights to those miRNAs also present in Gen3G resulted in a miR${}_{26}$Clock-GA with a MSE of 1.3 (RMSE=1.14). The correlation between chronological GA and miR${}_{26}$Clock-GA was 0.28 ($R^{2}$=0.08; shown in **Figure 6**).

As illustrated in **Figure 5** in the article, and **Supplementary Table 11**, the correlation between chronological GA and miRClock-GA was consistent across 10 different train–test splits, regardless of the type of lambda (minimal or 1se) applied. In the training sets, miRClock-GA derived using the minimal lambda showed substantially higher correlations with chronological GA compared to miRClock-GA derived using 1se lambda. However, in the test sets the differences between the minimal and 1se lambdas were smaller. Notably, minimal lambda still achieved slightly higher correlations in the test sets. Altogether the results suggest that the higher number of included miRNA in the miRClock-GA based on the minimal lambda provides better performance in unseen data. However, this gain was marginal compared tot he training data, indicating that the higher performance is offset to a large degree by a tendency for overfitting, resulting in an overall small net gain in performance in independent datasets.

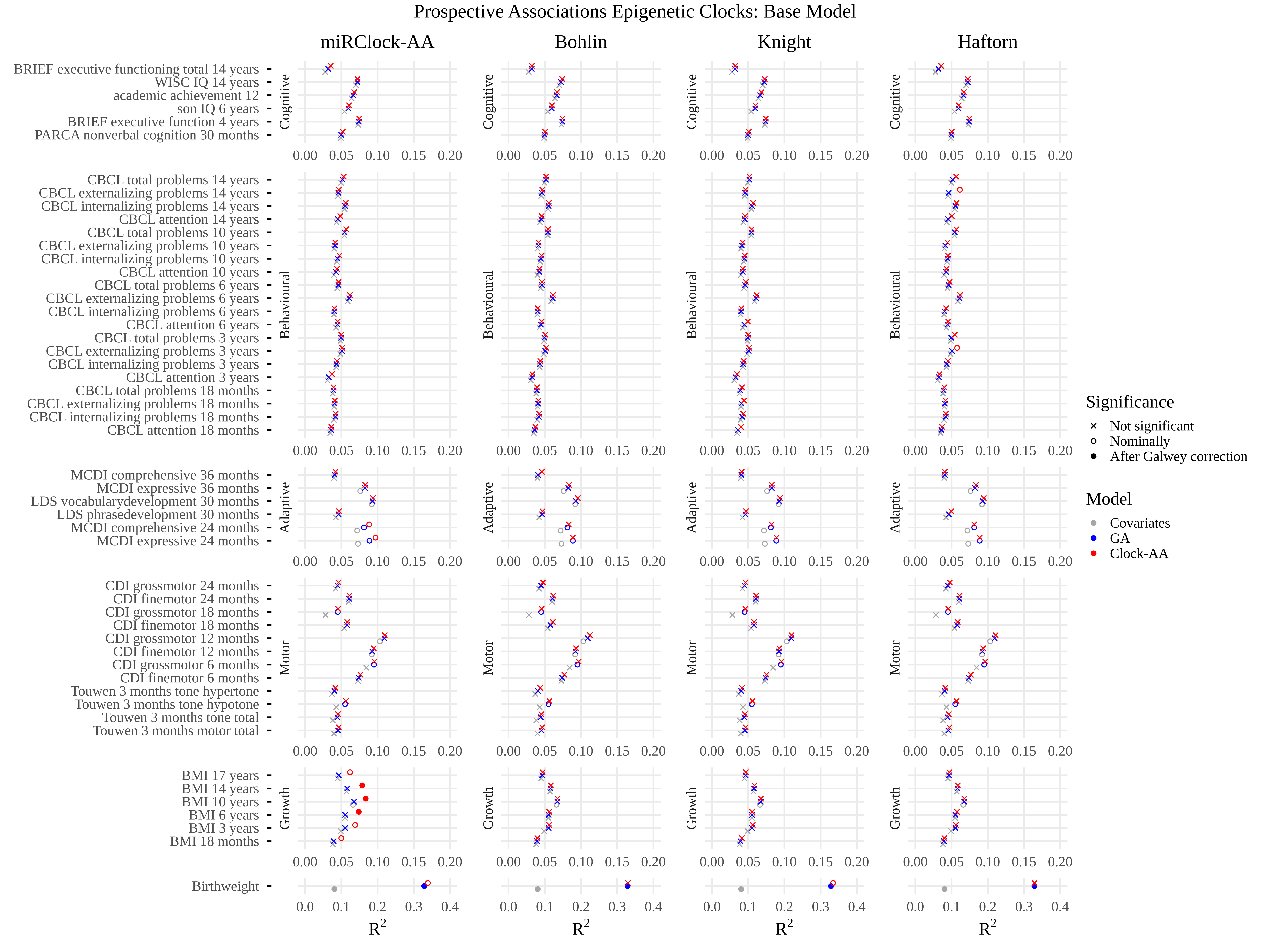

*Figure S1. Prospective associations between the epigenetic clocks (miRClock-AA, DNAmClock-AA) and behavioural and health-related phenotypes at later ages. These are the base models, which did not include corrections for maternal variables. The covariate models (in grey) cover cumulative variance explained by sex, haemolysis, miRNA-sequencing plate, initial total sample miRNA concentrations prior to dilution, and blood cell counts as estimated from DNAm data. The GA models (in blue) cover the covariate model in addition to clinical GA. The miRClock-AA models (in red) cover the GA model in addition to miRClock-AA. MiRClock-AA in this figure is derived from miRClock-GA based on the minimal lambda.*

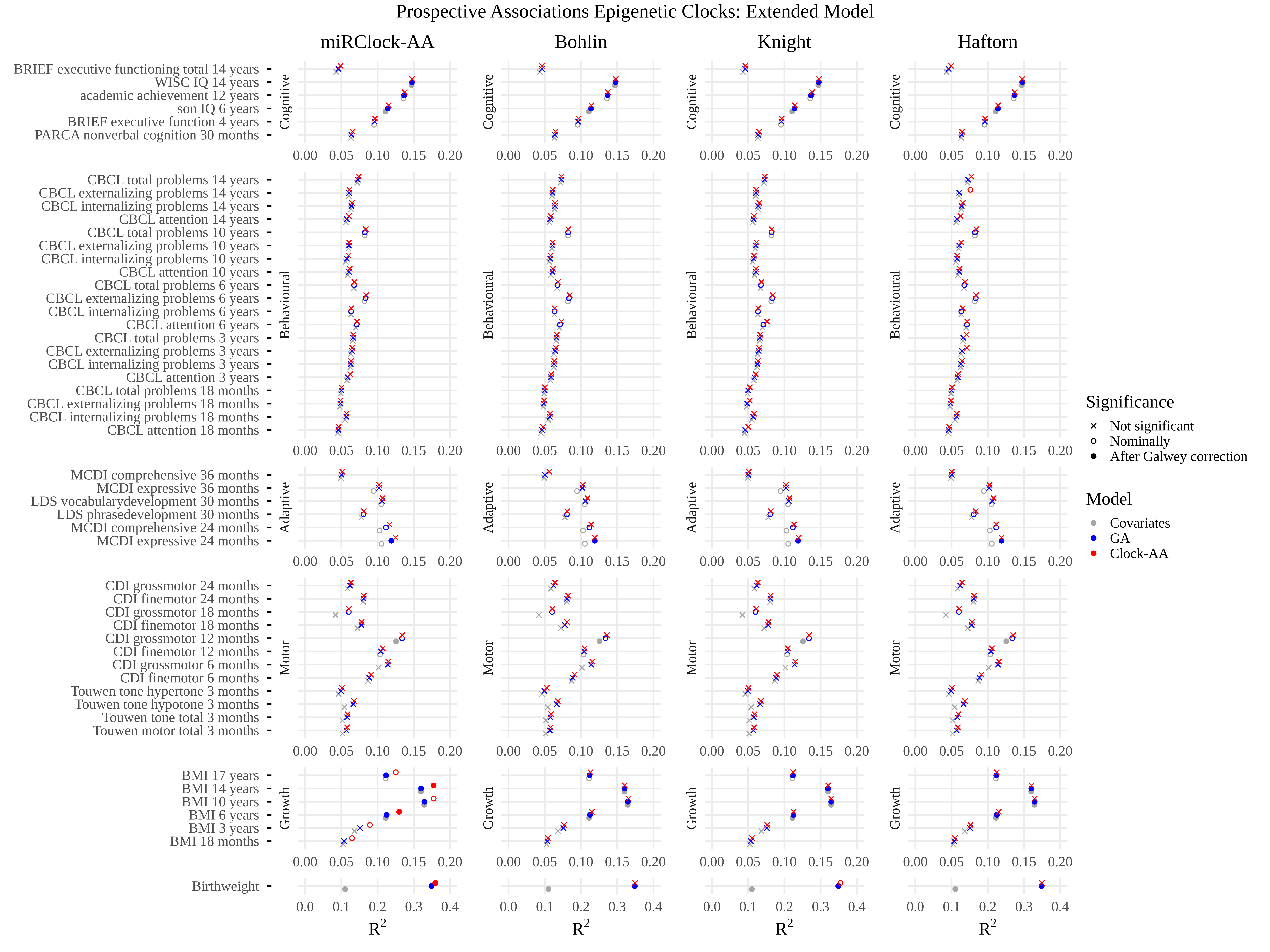

*Figure S2. Prospective associations between the epigenetic clocks (miRClock-AA, DNAmClock-AA) and behavioural and health-related phenotypes at later ages. These are the extended models, which included maternal covariates such as maternal age, education level, BMI, and smoking status. The covariate models (in grey) cover cumulative variance explained by sex, haemolysis, miRNA-sequencing plate, initial total sample miRNA concentrations prior to dilution, and blood cell counts as estimated from DNAm data. The GA models (in blue) cover the covariate model in addition to clinical GA. The miRClock-AA models (in red) cover the GA model in addition to miRClock-AA. MiRClock-AA in this figure is derived from miRClock-GA based on the minimal lambda.*

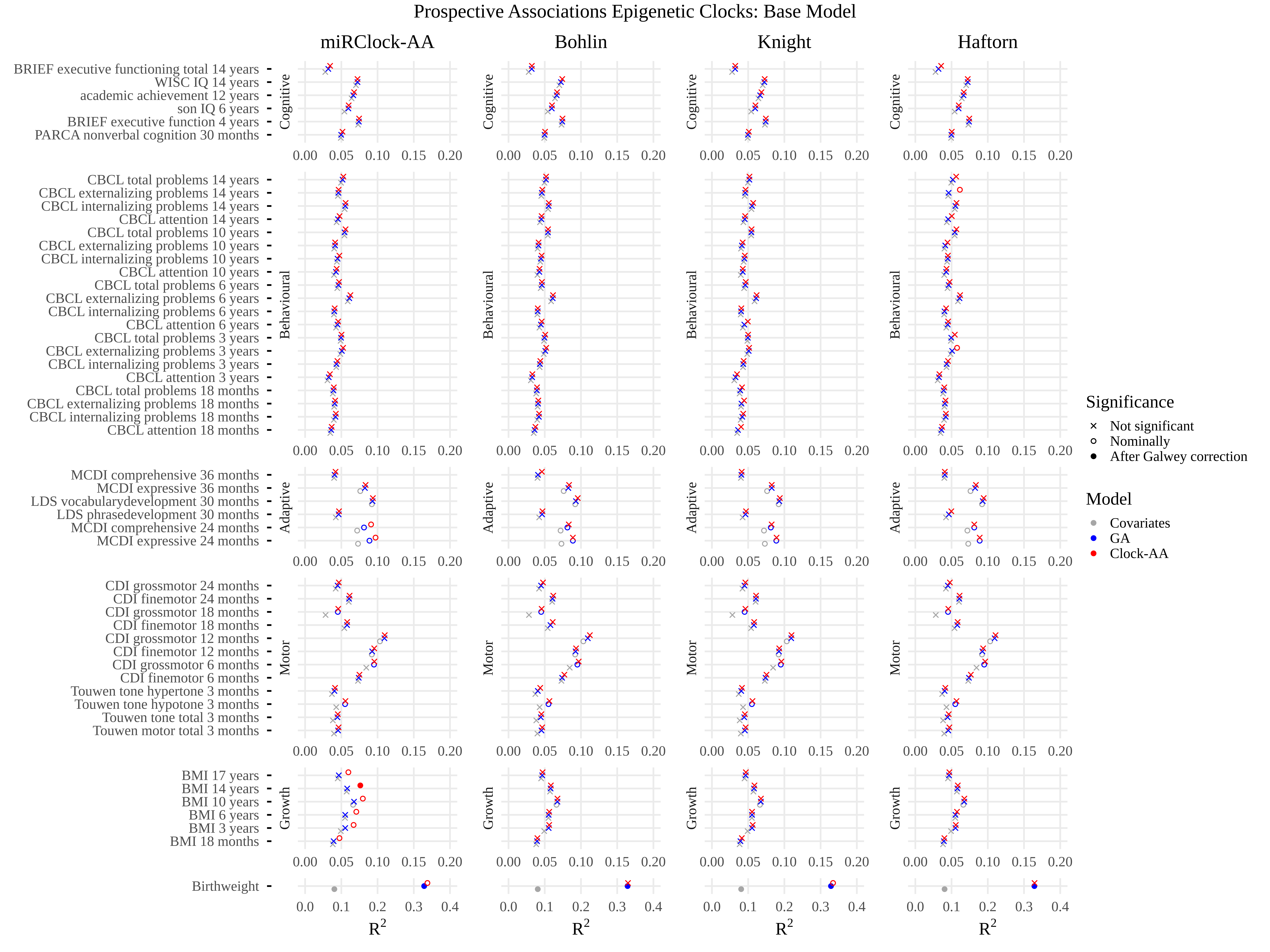

*Figure S3. Prospective associations between the epigenetic clocks (miRClock-AA, DNAmClock-AA) and behavioural and health-related phenotypes at later ages. These are the base models, which did not include corrections for maternal variables. The covariate models (in grey) cover cumulative variance explained by sex, haemolysis, miRNA-sequencing plate, initial total sample miRNA concentrations prior to dilution, and blood cell counts as estimated from DNAm data. The GA models (in blue) cover the covariate model in addition to clinical GA. The miRClock-AA models (in red) cover the GA model in addition to miRClock-AA. MiRClock-AA in this figure is derived from miRClock-GA based on the 1SE lambda.*

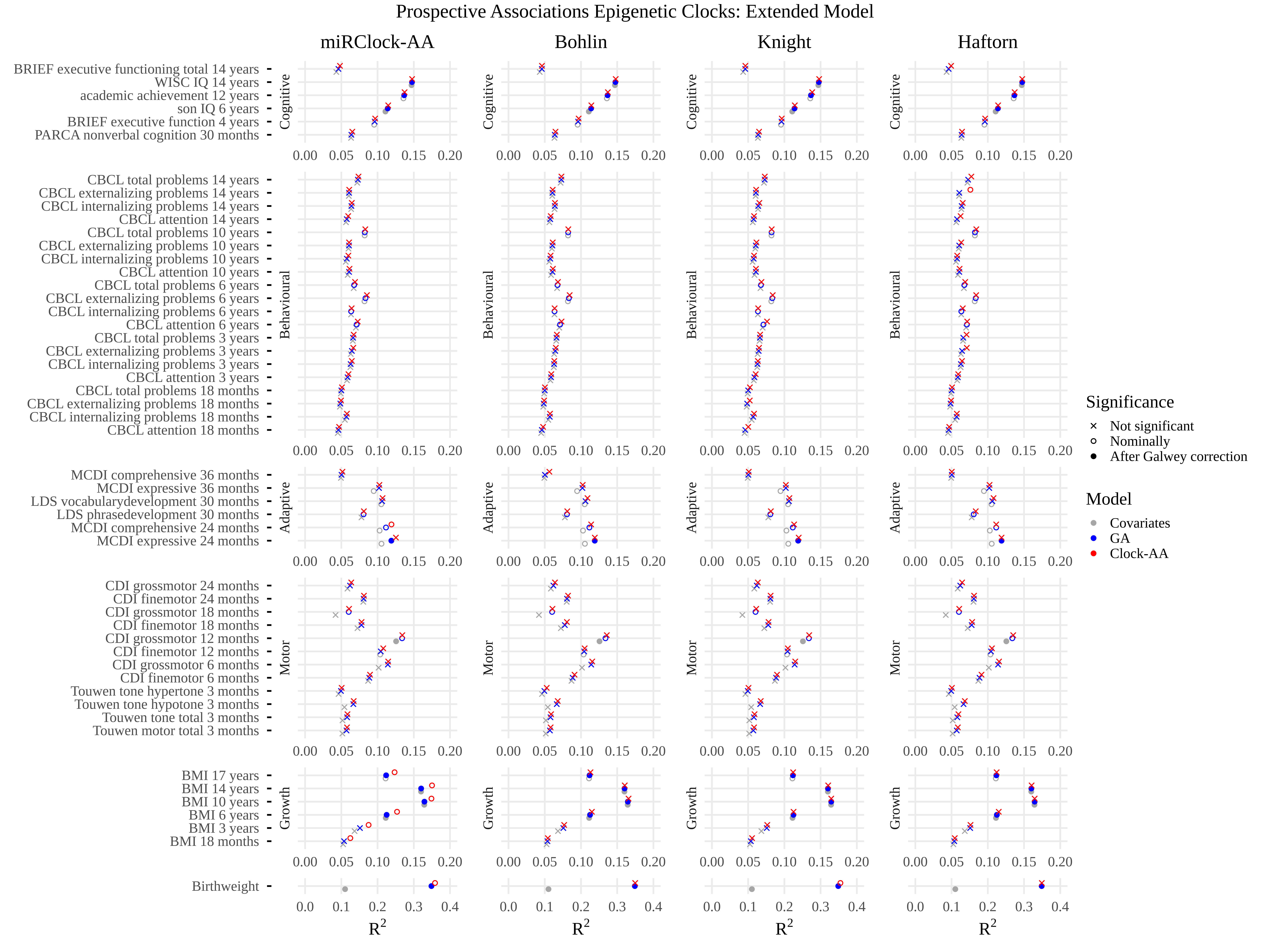

*Figure S4. Prospective associations between the epigenetic clocks (miRClock-AA, DNAmClock-AA) and behavioural and health-related phenotypes at later ages. These are the extended models, which included maternal covariates such as maternal age, education level, BMI, and smoking status. The covariate models (in grey) cover cumulative variance explained by sex, haemolysis, miRNA-sequencing plate, initial total sample miRNA concentrations prior to dilution, and blood cell counts as estimated from DNAm data. The GA models (in blue) cover the covariate model in addition to clinical GA. The miRClock-AA models (in red) cover the GA model in addition to miRClock-AA. MiRClock-AA in this figure is derived from miRClock-GA based on the 1SE lambda.*

##### Consistency of prospective associations between miRClock-GA and health and developmental outcomes over all train-test splits

The observed association between miRClock-GA and birthweight was replicated in all train-test splits, indicating a consistent relation. The association between miRClock-AA and birthweight remained significant after Galwey-correction for multiple testing in four out of ten base models (**Figure S5**), and in five out of ten extended models (**Figure S6**). Notable is that in three models, the association is Galwey-significant for both base and extended model, while in two base models a nominally significant association is followed up by Galwey-significance in the corresponding extended model, and the reverse is true for one model.

Other outcomes with Galwey-significant associations with miRClock-GA in the base model, were MCDI expressive language at 2 years old (in 2 out of 10 train-test splits), BMI at 10 years (in 1 out of 10), and BMI at 14 years (in 1 out of 10). For the extended model, substantially more outcomes showed Galwey-significant associations with miRClock-GA. These were: BMI at 10, 14, and 17 years, as well as WISC IQ at 14 years (all in 10 out of 10 train-test splits); BMI at 6 years and SON IQ at 6 years (both in 8 out of 10); academic achievement (in 7 out of 10); CBCL total problems at 10 years (in 6 out of 10); CBCL total problems at 6 years (in 4 out of 10); MCDI expressive language at 2 years (in 3 out of 10); CBCL attention problems at 6 years, CBCL externalizing problems at 6 years, and MCDI comprehensive language at 2 years (in 2 out of 10); and BMI at 2 years, BRIEF executive functioning total scale at 14 years, CBCL attention problems at 10 years, CBCL externalizing problems at 10 and at 14 years, CBCL internalizing problems at 6 years, CBCL total problems at 14 years, MCDI expressive language at 3 years (all in 1 out of 10).

Other outcomes with Galwey-significant associations with miRClock-AA in the base model (**Figure S1**), were BMI at 6, 10, and 14 years. These were all observed in 1 out of 10 train-test splits. For the extended model (**Figure S2**), BMI at 6 and 14 (both in 1 out of 10 train-test splits) showed Galwey-significant associations with miRClock-GA.

In the prospective associations between outcomes and miRClock-GA based on lambda 1se, we observe similar variations in achieving statistical significance over the different train-test sets (**Figure S7**; **Figure S8**).

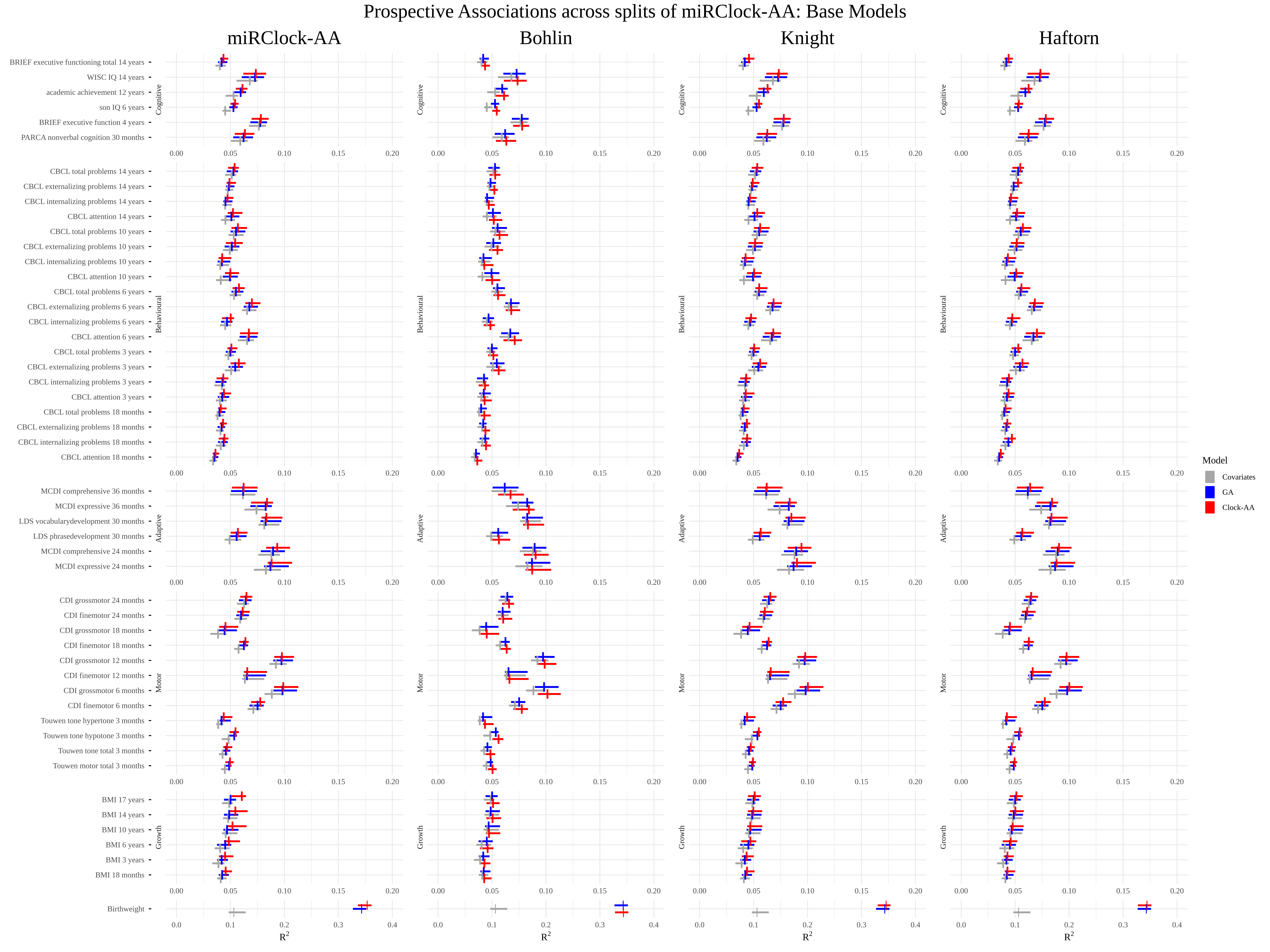

*Figure S5. Prospective associations between epigenetic clocks (miRClock-AA, DNAmClock-AA) and behavioural and health-related phenotypes at later ages across all splits of miRClock-GA test sets. These are the base models. The covariate models (in grey) cover cumulative variance explained by sex, haemolysis, miRNA-sequencing plate, initial total sample miRNA concentrations prior to dilution, and blood cell counts as estimated from DNAm data. The GA models (in blue) cover the covariate model in addition to clinical GA. The miRClock-AA models (in red) cover the GA model in addition to miRClock-AA. Vertical lines refer to the median value of the observed associations over all 10 train-test splits, while the horizontal lines refer to 95% confidence intervals.*

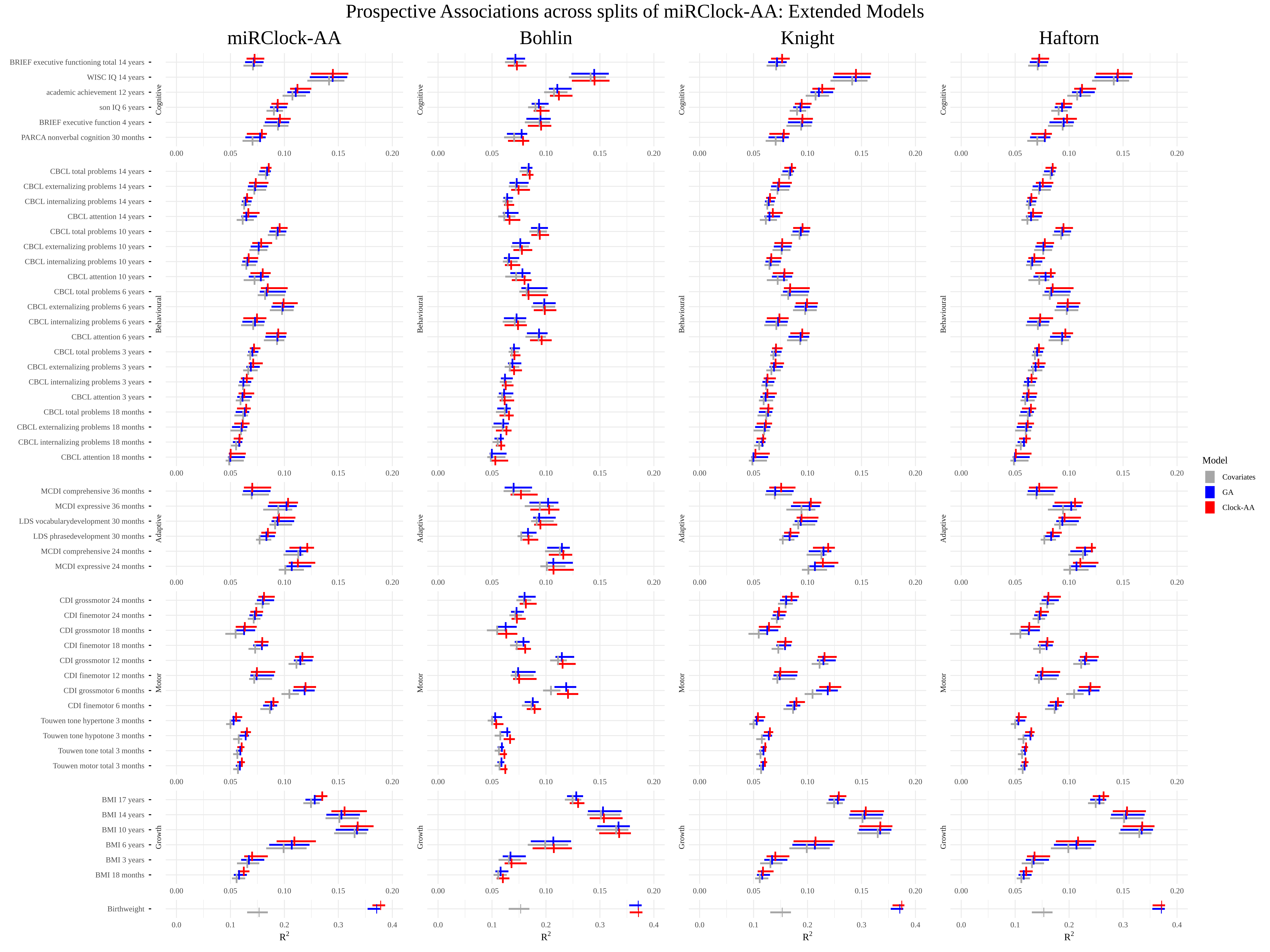

*Figure S6. Prospective associations between epigenetic clocks (miRClock-AA, DNAmClock-AA) and behavioural and health-related phenotypes at later ages across all splits of miRClock-GA test sets. These are the extended models, which included maternal covariates such as maternal age, education level, BMI, and smoking status. The covariate models (in grey) cover cumulative variance explained by sex, haemolysis, miRNA-sequencing plate, initial total sample miRNA concentrations prior to dilution, and blood cell counts as estimated from DNAm data. The GA models (in blue) cover the covariate model in addition to clinical GA. The miRClock-AA models (in red) cover the GA model in addition to miRClock-AA. Vertical lines refer to the median value of the observed associations over all 10 train-test splits, while the horizontal lines refer to 95% confidence intervals.*

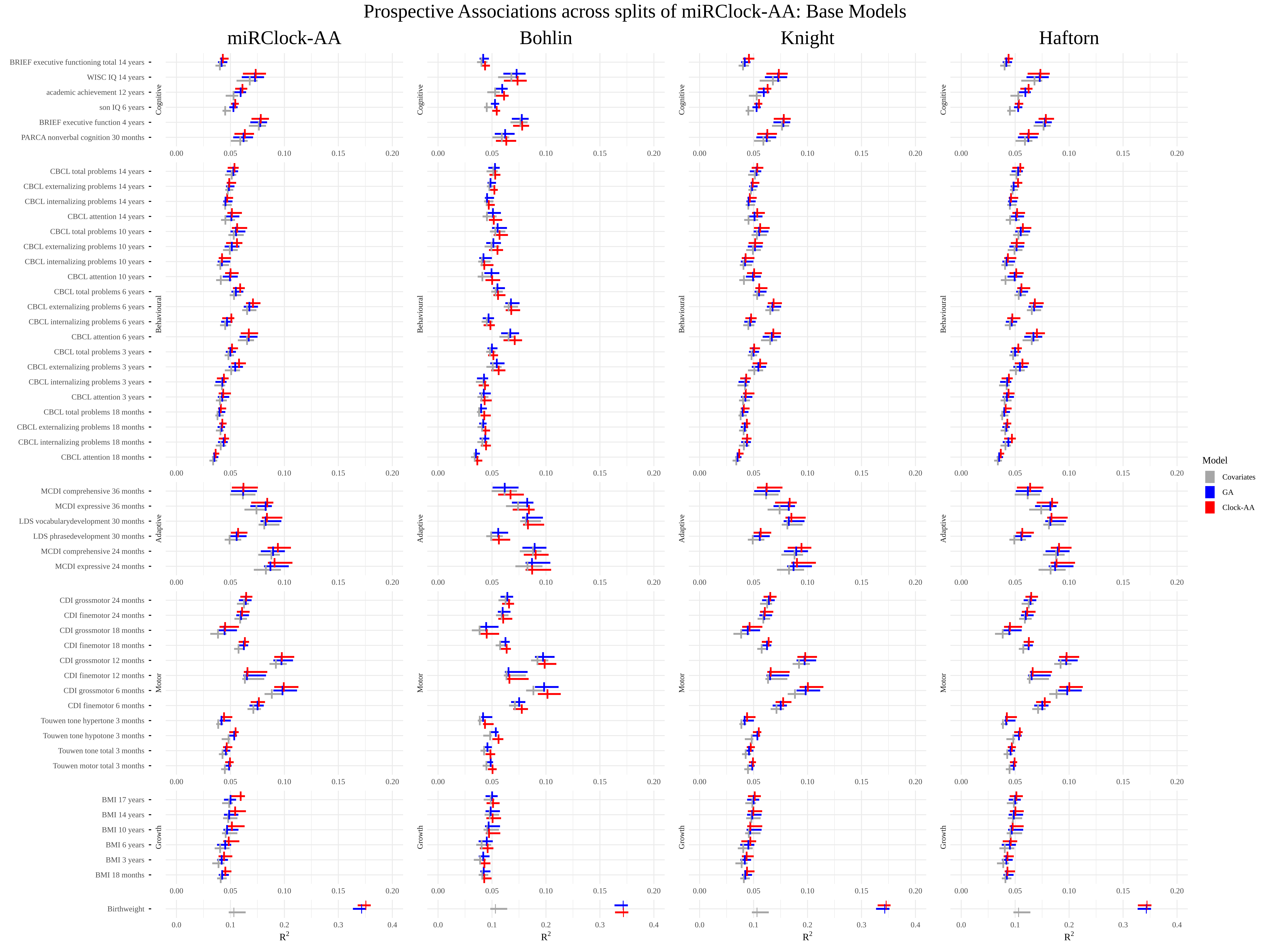

*Figure S7. Prospective associations between epigenetic clocks (miRClock-AA, DNAmClock-AA) and behavioural and health-related phenotypes at later ages across all splits of miRClock-GA test sets. These are the base models. The covariate models (in grey) cover cumulative variance explained by sex, haemolysis, miRNA-sequencing plate, initial total sample miRNA concentrations prior to dilution, and blood cell counts as estimated from DNAm data. The GA models (in blue) cover the covariate model in addition to clinical GA. The miRClock-AA models (in red) cover the GA model in addition to miRClock-AA. Vertical lines refer to the median value of the observed associations over all 10 train-test splits, while the horizontal lines refer to 95% confidence intervals.*

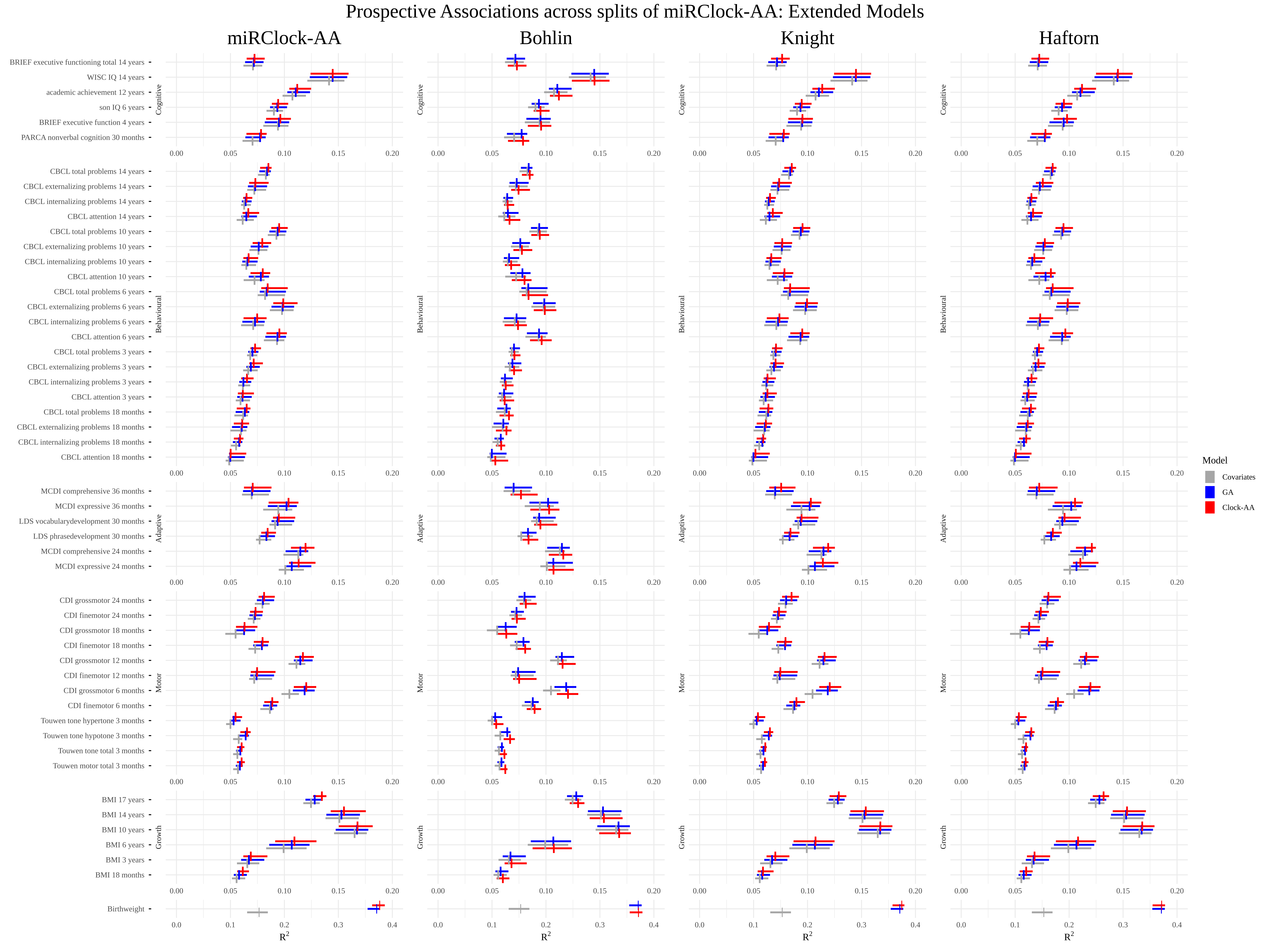

*Figure S8. Prospective associations between epigenetic clocks (miRClock-AA, DNAmClock-AA) and behavioural and health-related phenotypes at later ages across all splits of miRClock-GA test sets. These are the extended models, which included maternal covariates such as maternal age, education level, BMI, and smoking status. The covariate models (in grey) cover cumulative variance explained by sex, haemolysis, miRNA-sequencing plate, initial total sample miRNA concentrations prior to dilution, and blood cell counts as estimated from DNAm data. The GA models (in blue) cover the covariate model in addition to clinical GA. The miRClock-AA models (in red) cover the GA model in addition to miRClock-AA. Vertical lines refer to the median value of the observed associations over all 10 train-test splits, while the horizontal lines refer to 95% confidence intervals.*

##### Deviation of miRClock-GA from chronological GA with 1SE lambda

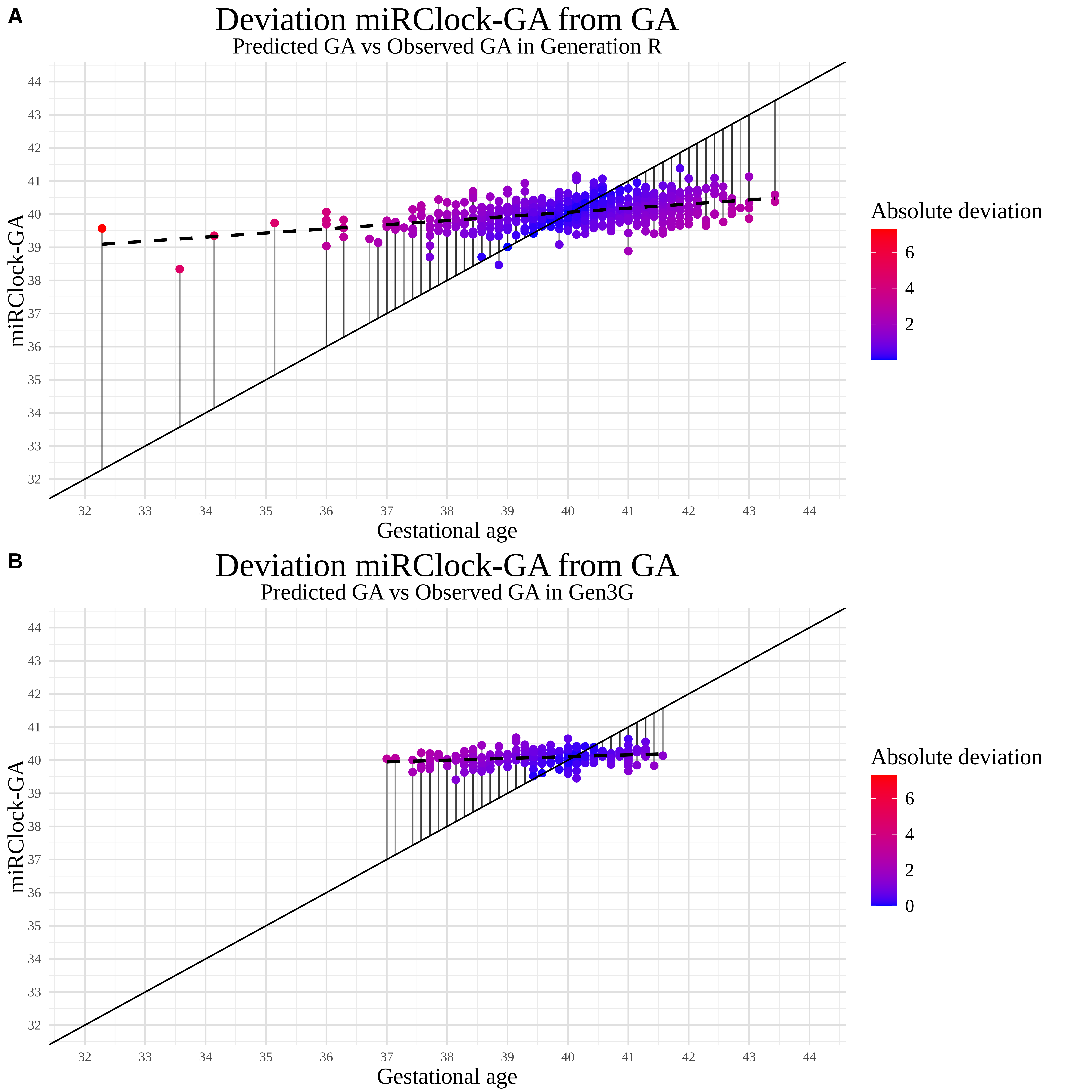

*Figure S9. Deviation of miRClock-GA (lambda 1se) relative to chronological GA for both (A) Generation R- and (B) Gen3G-cohorts. The solid line represents a hypothetical perfect predicted value of GA, with lines from each miRClock-GA value towards this line indicating the corresponding deviation. The colours of the dots indicate the degree of absolute deviation of miRClock-GA from the perfect prediction. The dotted line represents a regression line drawn based on regressing GA onto miRClock-GA.*

##### Sensitivity analysis: GA check on full GenR-cohort

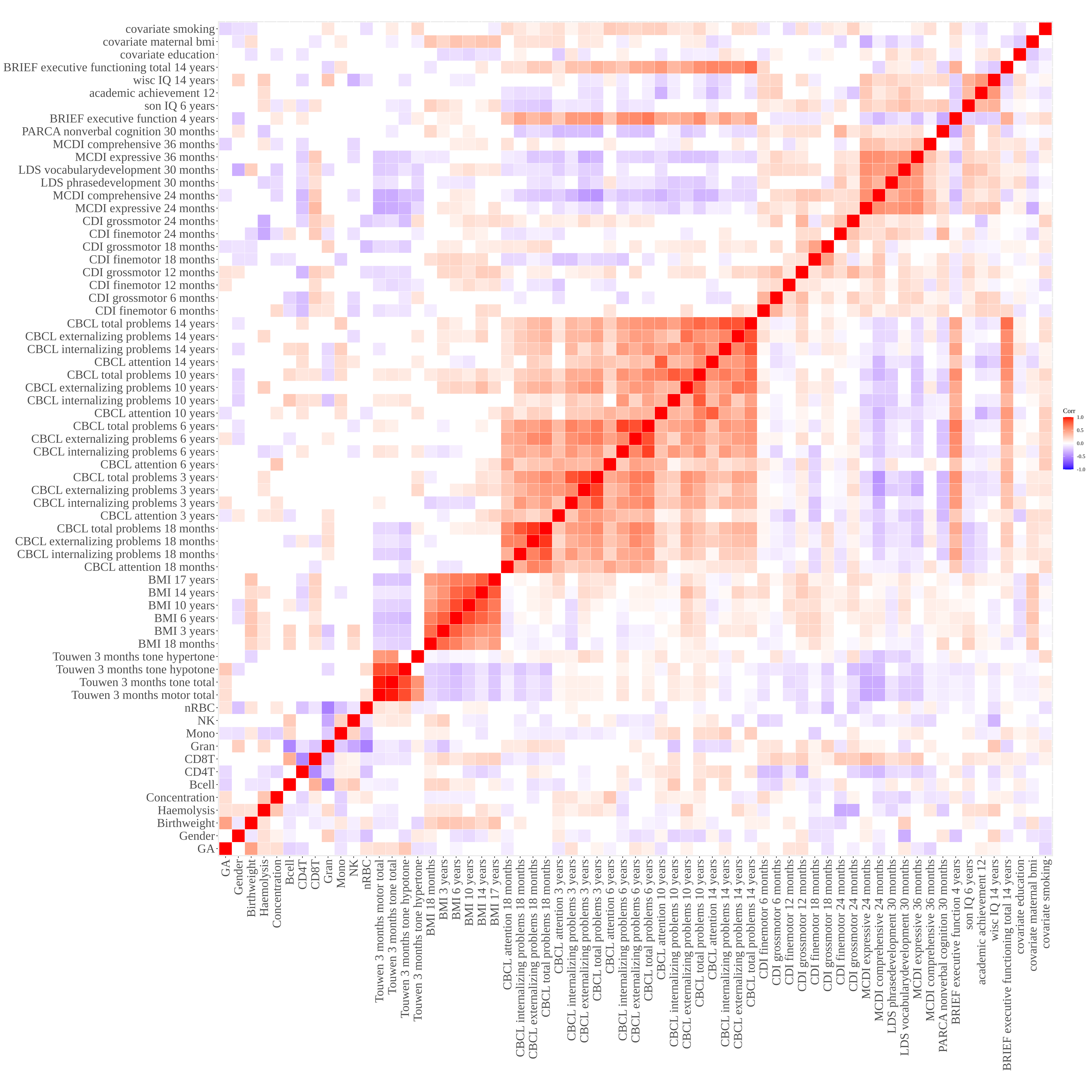

*Figure S10. Correlation matrix over the complete Generation R-cohort for chronological GA, covariates, and prospective outcomes included in Step 3.*
